# A generic noninvasive neuromotor interface for human-computer interaction

**DOI:** 10.1101/2024.02.23.581779

**Authors:** CTRL-labs at Reality Labs, David Sussillo, Patrick Kaifosh, Thomas Reardon

## Abstract

Since the advent of computing, humans have sought computer input technologies that are expressive, intuitive, and universal. While diverse modalities have been developed, including keyboards, mice, and touchscreens, they require interaction with an intermediary device that can be limiting, especially in mobile scenarios. Gesture-based systems utilize cameras or inertial sensors to avoid an intermediary device, but they tend to perform well only for unobscured or overt movements. Brain computer interfaces (BCIs) have been imagined for decades to solve the interface problem by allowing for input to computers via thought alone. However high-bandwidth communication has only been demonstrated using invasive BCIs with decoders designed for single individuals, and so cannot scale to the general public. In contrast, neuromotor signals found at the muscle offer access to subtle gestures and force information. Here we describe the development of a noninvasive neuromotor interface that allows for computer input using surface electromyography (sEMG). We developed a highly-sensitive and robust hardware platform that is easily donned/doffed to sense myoelectric activity at the wrist and transform intentional neuromotor commands into computer input. We paired this device with an infrastructure optimized to collect training data from thousands of consenting participants, which allowed us to develop generic sEMG neural network decoding models that work across many people without the need for per-person calibration. Test users not included in the training set demonstrate closed-loop median performance of gesture decoding at 0.5 target acquisitions per second in a continuous navigation task, 0.9 gesture detections per second in a discrete gesture task, and handwriting at 17.0 adjusted words per minute. We demonstrate that input bandwidth can be further improved up to 30% by personalizing sEMG decoding models to the individual, anticipating a future in which humans and machines co-adapt to provide seamless translation of human intent. To our knowledge this is the first high-bandwidth neuromotor interface that directly leverages biosignals with performant out-of-the-box generalization across people.

## Introduction

Interactions with computers are increasingly ubiquitous, but existing input modalities suffer from persistent tradeoffs between portability, throughput, and accessibility. The concept of a neural interface that can obviate these tradeoffs and provide seamless interaction between humans and machines at the speed of thought has long been a staple of science fiction, but has been slower to emerge in reality. In recent years, intracortical neural interfaces that directly interface with brain tissue have advanced the premise (Gilja et al. 2012; Hochberg et al. 2012), demonstrating translation of thought into language at bandwidth rates comparable with conventional computer input systems (Collinger et al. 2013; Willett et al. 2023). However, existing high-bandwidth interfaces rely on invasive neurosurgery, and the underlying models that translate neural signals to digital inputs remain bespoke. Non-invasive approaches relying on recording of electroencephalogram (EEG) (Abiri et al. 2019) signals at the scalp have offered more generality across people, e.g., for entertainment (Kerous, Skola, and Liarokapis 2018). But EEG can require lengthy setup, and the problematic signal-to-noise ratio (SNR) of these devices has limited their utility (Défossez et al. 2023). Regardless of the modality, issues of signal bandwidth, generalization across populations, and the desire to avoid per-person or session-to-session calibration remain key technical hurdles in the field of brain computer interfaces (BCI) (Brandman, Cash, and Hochberg 2017; Lotte et al. 2018; Gilja et al. 2012; Degenhart et al. 2020; Brandman et al. 2018).

To build an interface that is both performant and accessible, we focused on an alternate class of non-invasive *neuromotor interfaces*, which are based on reading out the electrical signals from muscles, i.e. electromyography (EMG). Myoelectric potentials are produced by the summation of motor unit action potentials (MUAPs) and thus represent a window into the motor commands issued by the central nervous system. Surface EMG (sEMG) recordings offer high SNR by virtue of amplification of neural signals in the muscle (Kandel et al. 2000), enabling real-time single-trial gesture decoding. Further, sEMG does not suffer from problems that vex computer-vision-based human-computer interaction (HCI) approaches, such as occlusion, lighting, or gestures with minimal movement. EMG is widely used in the clinic, for example, to study muscle activation (De Luca 1997), diagnose disorders (Pullman et al. 2000), and foster and monitor neurorehabilitation (Farina et al. 2014; Campanini et al. 2020; Merletti and Farina 2016). It has also been used in human-machine interfaces dating back to the 1950s (Battye, Nightingale, and Whillis 1955; Berger and Huppert 1952).

Current EMG systems have many limitations for wide-scale use and deployment. Most are limited to in-lab and supervised research applications while multi-channel sEMG setups are generally encumbered with wires to external amplifiers and power sources. Many sEMG systems are placed over unconventional or uncomfortable locations such as the target muscle’s belly to ensure high signal quality (Merletti and Muceli 2019; Del Vecchio et al. 2020). Even commercially available EMG-based neuromotor interfaces have been historically challenging to control (Biddiss and Chau 2007), relating to myriad technical issues such as poor robustness across postures (Scheme et al. 2010), lack of standardized data (Phinyomark and Scheme 2018), electrode displacement (Young, Hargrove, and Kuiken 2011), and lack of both cross-session (Zia Ur Rehman et al. 2018) and cross-user generalization (Saponas et al. 2008). More recently, deep learning techniques have shown some success at addressing these limitations, e.g., (Côté-Allard et al. 2019), but a general lack of available EMG data and low sample sizes are believed to limit their efficacy (Phinyomark and Scheme 2018). In summary, while sEMG-based approaches show promise due to high SNR, the question of whether they can be applied to a large population in practical settings to solve practical HCI problems remains open.

To validate the thesis that sEMG can provide an intuitive and seamless computer input that works in practice across a population, we developed and deployed robust, non-invasive hardware for recording sEMG at the wrist. We chose the wrist because humans primarily engage the world with their hands, and the wrist provides broad coverage of sEMG signals of hand, wrist, and forearm muscles while affording social acceptability for wireless wearable technology, such as smart watches (Jiang et al. 2018; Mendez Guerra et al. 2022). Our sEMG research device (sEMG-RD) is a dry-electrode, multi-channel recording platform with much higher sample rates (2kHz) than past consumer sEMG devices (Mendez et al. 2017), and is capable of extracting single putative MUAPs. It is comfortable, wireless, accommodates diverse anatomy and environments, and can be donned/doffed in a few seconds.

To transform sEMG into commands that could drive computer interactions, we architected and deployed neural networks trained on data from thousands of consenting human participants (e.g. >6,500 participants for one network, see Online Methods). We also created automated behavioral prompting and participant selection systems to scale neuromotor recordings across a large and diverse population. We demonstrated the ability of our sEMG-RD to drive different computer interactions such as 1D continuous navigation (akin to pointing a laser-pointer based on wrist posture), gesture detection (finger pinches and thumb swipes), and handwriting transcription. The sEMG decoding models performed well across many people without person-specific training or calibration, including across-session recalibration. In offline evaluation, our sEMG-RD platform achieved greater than 90% classification accuracy for held-out participants in handwriting and gesture detection, as well as greater than 75% accuracy on wrist pose classification. We achieved 0.5 target acquisitions per second in wrist pose-based continuous navigation, 0.9 acquisitions per second on discrete gestures, and 17.0 adjusted words per minute (aWPM) with handwriting on computer-based tasks that evaluate these interactions in closed-loop.

Concretely, our scientific innovations include the following. We developed dry electrode sEMG hardware with high SNR that is easily donned/doffed and that accommodates diverse human anatomy. Enabled by this hardware platform, we performed data collection at scale to train large neural network models on thousands of users. We showed that these networks are capable of generalizing with high performance to new users in closed loop settings, thereby demonstrating that sEMG serves as a viable biosignal for addressing unsolved HCI problems at scale. We characterized the models’ performance scaling with respect to the number of users in the dataset and provide a robust analysis of the effects of personalization.

To our knowledge, this is the first neuromotor interface that achieves such a high level of performance and scalability. Our sEMG-RD platform has demonstrated the ability to drive various computer interactions with high accuracy across a large and diverse population, without the need for person-specific training or calibration. Our approach opens up new directions of EMG-based HCI research and development while simultaneously solving many of the technical problems that are of fundamental importance to current and future BCI efforts, whether intracortical (Willett et al. 2023; Musk and Neuralink 2019; Metzger et al. 2023) or non-invasive.

## Results

### Scalable electromyography recording platform

To build generic data-driven sEMG decoding models capable of predicting user intent from neuromuscular signals, we developed a hardware and software platform capable of robustly interfacing the neuromotor system with computers with minimal setup time across a diverse population (Fig. 1a). Consenting participants (see Online Methods for participant onboarding and consent) were seated in front of a computer while wearing an sEMG research device (sEMG-RD) at the wrist: a dry electrode, multi-channel recording platform with high sample rate (2 kHz), and low-noise (2.46 µVrms) compatible with everyday use (Fig. 1a) (Jiang et al. 2018; Mendez Guerra et al. 2022) (Online Methods). We fabricated the device in 4 different sizes to ensure coverage across a wide range of wrist circumferences. The device streamed wirelessly over secure Bluetooth standard protocols and provided a battery life of more than 4 hours.

**Fig. 1.**
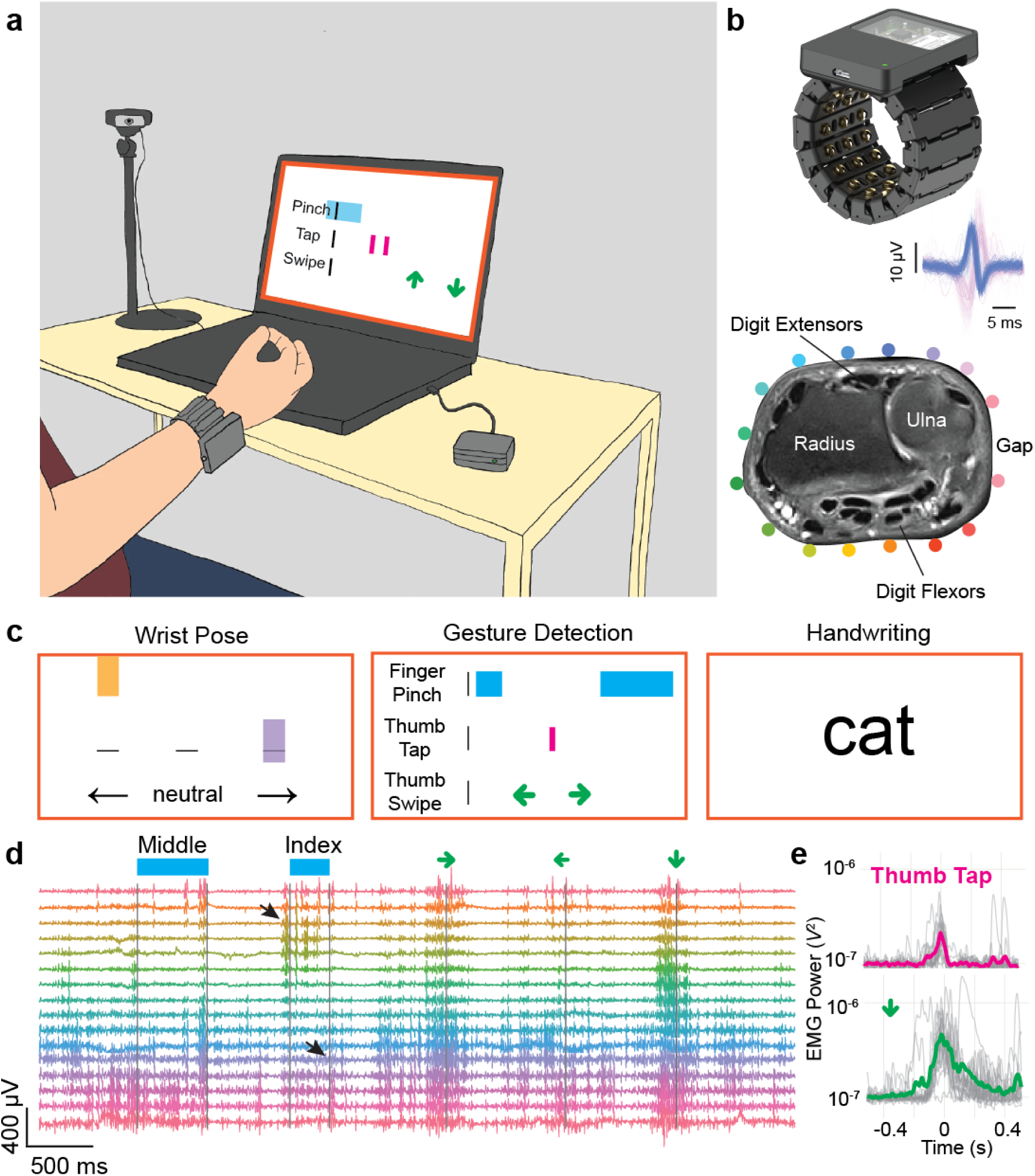
A hardware and software platform for high-throughput, real-time surface EMG decoding at the wrist. **a**, Overview of sEMG data collection and computer interaction. A participant wears the sEMG wristband which communicates to a computer using a Bluetooth receiver. The participant is prompted on a computer screen to perform movements using programmatically generated prompts. A webcam captures the participant’s hand and wrist while excluding the face from the field of view. During different experimental protocols participants are instructed to perform diverse movements of the hand and wrist: deflecting the wrist to different cardinal directions, performing discrete gestures, and handwriting of text. Between different sessions participants are instructed to remove and slightly reposition the band to enable generalization across different recording positions. **b**, (*Upper*) The sEMG wrist band consists of 48 electrode pins configured into 16 bipolar channels along the proximal-distal axis of the forearm and the remainder serving as shield and ground. A 3D printed housing encloses cabling and analog amplifiers for each channel. A computing core digitizes the signal and streams sEMG data using Bluetooth. (*Inset*) Overlay of 62 and 72 individual instances of two putative MUAPs evoked by subtle thumb extension (blue) and pinky extension (pink) movements, respectively. Shown are recordings from a single sEMG channel selected for having high variance during the movement (Online Methods). (*Lower*) A proton density weighted, axial plane MRI scan of the wrist, with relevant bone and muscle landmarks labeled. Colored dots indicate approximate position of electrodes, with an adjustable gap between electrodes placed over an area of low muscle density at the ulna. **c**, Schematic examples of prompters for the wrist pose (left), gesture detection (middle), and handwriting tasks (right). During each task, prompts move from top to bottom, left to right, or vary from screen to screen respectively, to instruct participants (see Extended Data Fig. 3). **d**, sEMG signals recorded during performance of discrete gestures and high-pass filtered at 20 Hz reveal intricate patterns of activity across multiple channels accompanying performance of each gesture, with prompt timings above (e.g. “Middle” indicates the timing of a middle pinch prompt, while a green left arrow indicates the same for a leftward thumb swipe). Channel coloring corresponds to electrode locations in b. Black arrowheads highlight activation of flexors during an index-to-thumb pinch and extensors during the release of the index finger from thumb contact. **e**, Variability in gestural sEMG activations across instances (top shows thumb taps, bottom shows downward thumb swipe). The gray lines show the instantaneous high-pass filtered sEMG power, summed across channels, for all instances of a gesture during a single band placement. The bold traces show the average. The mean was subtracted from all traces, and the power was offset by 10^−7^ *V*^2^ to plot on a logarithmic scale without visually exaggerating baseline variance.

We optimized the sEMG-RD design for recording subtle electrical potentials at the wrist. The device used 48 gold-plated electrodes configured into 16 bipolar EMG-sensing channels oriented along the proximo-distal axis of the forearm and the remainder serving as shield and ground (Extended Data Fig. 1). We chose the circumferential inter-electrode spacing of 10.6–15 mm, varying with the device size, to approach the spatial bandwidth of EMG signals at the forearm (∼5-10 mm) (Merletti and Farina 2016), while minimizing the device’s form factor. We placed the gap in electrodes to allow for tightening adjustments along the ulna bone, where muscles are reduced in density. Together, this allowed for sensing putative MUAPs across the wrist (Fig. 1b)

To collect training data for models, we recruited an anthropometrically and demographically diverse group of participants (200-6350 participants, depending on task; Extended Data Fig. 2) to perform three different tasks: continuous navigation, discrete gesture recognition, and handwriting. In all cases, participants wore sEMG bands on their dominant hand and were prompted to perform actions using a custom prompting system, run on laptops (Fig. 1c). For continuous navigation, participants were prompted to flex, extend, or deviate their wrist. During the discrete gesture detection task, a prompter instructed participants to perform nine distinct gestures with randomized gesture order and inter-gesture interval. During the handwriting task, participants were prompted to hold their fingers together (as if holding an imaginary writing implement) and “write” the prompted text. See Online Methods for further training data protocol details.

We designed the data collection system to facilitate supervised training of sEMG decoding models. During data collection, we recorded both sEMG activity and the timestamps of labels on the prompter using a real-time processing engine. We designed the engine to be used during recording and model inference to reduce online-offline shift (Online Methods). To precisely align prompter labels to actual gesture times, which may vary due to a participant’s reaction time or compliance, we developed a time alignment algorithm that allowed for *post hoc* inference of gesture event times (Online Methods).

Examination of raw sEMG traces revealed highly structured patterns of activity (Fig. 1d). During the gesture detection task, each action evoked patterned activity across a set of channels that roughly correspond to the position of flexor and extensor muscles for the corresponding movement (Fig. 1d, Extended Data Fig. 1c). The high sensitivity of the sEMG-RD allowed for observation of fine differences in sEMG power across instances of a given gesture performed during a session (Fig. 1e). This simultaneously highlights the power of the platform in acquiring repeated time-aligned examples for supervised learning and the challenges facing generalization of EMG decoders.

### Single-participant sEMG models do not generalize across individuals

It is well-known across BCI modalities that both across-session and across-user generalization are difficult problems (Gilja et al. 2012; Sussillo et al. 2016; Degenhart et al. 2020; Saponas et al. 2008). We wanted to evaluate the difficulty of these generalizations for sEMG decoders. Inspection of the raw data revealed pronounced variability in the sEMG for the same action across different participants and band donnings (which we refer to as sessions), reflective of variations in sensor placement, anatomy, physiology, and behavior that can make generalization challenging (Fig. 2a,b). The distribution of the distances between the event triggered average sEMG waveforms for a single gesture across different participants heavily overlapped with the distribution of distances across gestures, highlighting the challenge of the generalization problem (Fig. 2b, Extended Data Fig. 4a).

**Fig. 2.**
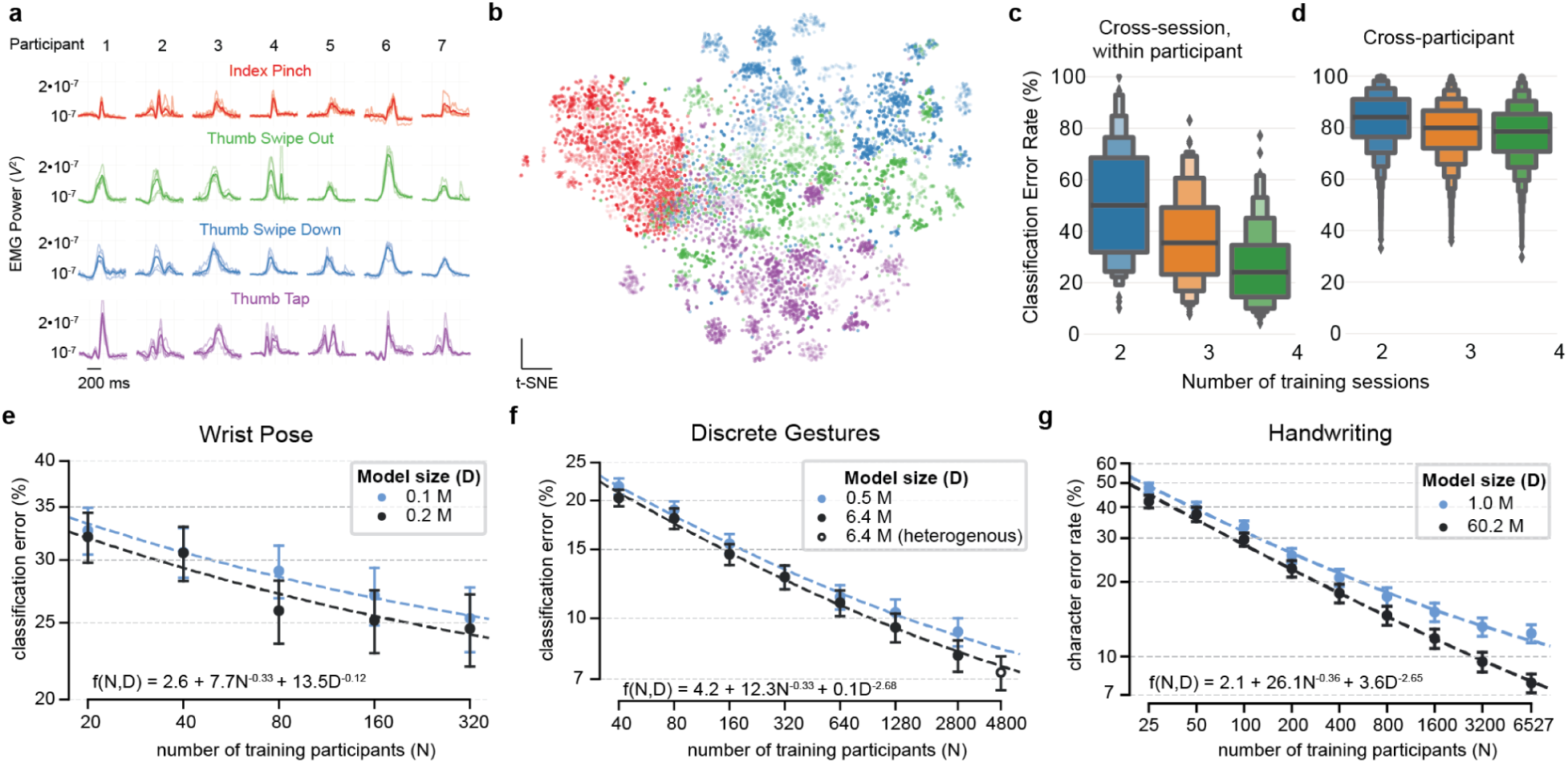
Generalization performance of single- and multi-participant models. a, Cross-participant (horizontal) and cross-session variability (light lines) in gestural sEMG for four discrete gestures (different rows and colors) across seven participants. Light lines are the high-pass filtered sEMG power, averaged across all channels and all gesture instances during a single band placement. The bold lines correspond to the average across all band placements. b, t-SNE embedding of sEMG activations across participants for the four different gestures in a and 20 participants. Colormap as in (a), with shading reflecting different participants. We used a concatenation of the 0-, 1- and 2-diagonals of the sEMG covariance matrix over a 300 ms window centered on each gesture instance to yield a 48×60-dimensional feature space (see Online Methods). t-SNE was run in 2 dimensions with perplexity 35 on the flattened feature space. c, Single-participant models trained and tested on the same participant show better generalization across sessions on discrete gesture classification as more training data is used. All pairwise comparisons are significant, *p* < 10^−10^, Wilcoxon signed-rank test. d, Models trained on a single participant generalize poorly across participants on discrete gesture classification even when more training data is used. All pairwise comparisons are significant, *p* < 10^−10^, Wilcoxon signed-rank test. e-g, Classification error of decoding models trained to classify (e) wrist poses (flexion, extension, neutral), (f) discrete gestures (up/down/left/right thumb swipes, thumb tap, index/middle press/release), and (g) handwritten characters (`a-z`, `0-9`, `,.?’!`, `space`), as a function of number of participants in the training set. Plots show mean +/− SEM classification error evaluated on a test set of held-out participants (39 for wrist pose, 100 for discrete gestures and 50 for handwriting). Inset equations show the form and parameters of the fitted scaling curves, plotted with dashed lines for each task and model size (N measured in units of 100’s of participants and D measured in millions of parameters). Note for the largest model for discrete gestures, indicated by an open circle, we used varying numbers of sessions per participant (see Online Methods for full details).

To evaluate the ability of obtaining performant sEMG decoders across sessions for a given participant, we trained single-participant models for 100 participants who had collected at least five sessions on a discrete gesture classification task with nine different gestures (press/release of index or middle finger to thumb, and thumb-based control: left/right/up/down/tap). For each participant, we held out one session for evaluation and then trained models on two, three, or four of the remaining sessions (see Online Methods for more details). As an offline evaluation metric we used the Classification Error Rate (CLER), defined as: one minus the number of correct predictions divided by the total number of true events.

Single-participant models trained and tested on the same participant achieved offline performance that improved substantially with more training data (Fig. 2c, from median CLER 50.6% ± 21.7% SEM with two training sessions to median CLER 26.5% ± 15.3% SEM with all 4 training sessions). In comparison, models trained on one participant and then tested on another showed substantially worse performance and benefited only mildly from an increasing amount of training data (Fig. 2d, from median CLER 82.9% ± 10.7% SEM with two training sessions to median CLER 77.7% ± 11.1% SEM with all 4 training sessions). The greater tractability of generalization across sessions versus generalization across people implies a significant domain shift across people. For most people, the model trained on their own data performed better compared to all other single-participant models (Extended Data Fig. 5a), and for all 100 people, their own model was within the top 5 performing models (Extended Data Fig. 5a).

We wondered whether cross-participant variability was difficult because there was structure or clusters across people, or whether every participant required a relatively unique single-participant model. The former could motivate an approach where a set of models trained on a small population (within each cluster) could achieve a high level of population coverage. Extended Data Fig. 5b shows a two-dimensional embedding using t-SNE with the distance between two participants being the average of the model transfer CLER between them. The absence of overt structure suggested that there are no immediately clear clusters of participants with quantitatively distinct performance statistics. Additionally, there are no people who exhibit a capacity to generate performant models for other people, nor are there any people for whom other people’s models always perform well. Extended Data Fig. 5c shows a scatter plot of the receiver score (CLER for a specific participant when using the models of all other participants) vs the donor score (CLER for each participant when using the model of a specific participant). The two metrics are weakly correlated and show high error rates (CLER > 60% in all cases).

### Ofline evaluation of generic models

To avoid the need to train and tune models for each individual, we trained *generic* models that are able to generalize across people. To do this, we collected large quantities of data from thousands of data collection participants. For each task (i.e. wrist pose, discrete gestures, and handwriting), data was collected open-loop to train the decoding models. See Online Methods for further details.

A natural question when training generic models is how performance of the model scales with the number of participants whose data is included in the training corpus. Of secondary interest is how this performance scaling relationship interacts with size of the architecture (i.e. number of model parameters). To explore these relationships, we looked at offline decoding performance of models trained on varying quantities of training data (Fig. 2e-g). The offline evaluation was performed entirely on held-out participants as a way of estimating model performance for a participant whose data was not included in the training set. The function fit to the empirical performance data is an inverse power law both as a function of parameters and data quantity, consistent with the scaling properties of large language models (Hoffmann et al. 2022) and vision transformers (Zhai et al. 2022). Note that the parameters of the scaling relationship are shared across architecture sizes (see Online Methods). In general we observed reliable performance improvements as a function of increasing number of participants in the training corpus as well as overall better performance for larger models.

### Online evaluation of generic models

Ultimately, closed-loop performance of our sEMG decoding models is the main metric that confirms their HCI utility. For each task, closed-loop evaluation was performed on naive participants who had not previously had meaningful experience using any sEMG decoder on these tasks (the same *N*=20 for wrist pose and handwriting, *N*=24 separate participants for discrete gestures). The core tasks were using the wrist pose decoder to continuously control a 1D cursor to acquire targets, the discrete gestures decoder to navigate and perform actions in a discrete lattice, and the handwriting decoder to write out prompted phrases that were then visualized on the screen (see Online Methods for evaluation tasks; see Supplementary Video 1, 2, 3, 4 for representative performance; see Extended Data Fig. 6 for depiction of task dynamics). For each task, participants performed 3 distinct blocks of trials to allow for characterization of learning, with the first block always being a practice block (10 trials for discrete gestures and handwriting, 50 trials for wrist pose) that allowed them to adapt to the controller.

For all tasks, we observed learning effects, where participants improved with experience. During the practice block, the study supervisor gave verbal coaching as needed to participants to encourage them to learn the gestures to complete each task. Coaching was along the lines of “hold your hand like this”, “perform the swipe faster”, “write more continuously, without pausing between each character”, etc. Coaching was actively provided by default only during the practice block, whereas in the evaluation blocks it was provided only on trials that users struggled to complete. The same style of coaching was used for evaluation blocks as during practice. While participants were typically able to perform each task on their own after the initial practice block, for the discrete gestures and handwriting tasks we found that coaching during the evaluation block was valuable for a subset of participants.

Every participant was able to complete every trial of the three tasks. For wrist pose control, all participants were able to successfully move the cursor to each target and stay on the targets for 500 ms to acquire them. Performance was characterized by time to target acquisition (Fig. 3d) and dial-in time, which measures the time taken to get back to the target after exiting it prematurely (Fig. 3e; see Online Methods for definitions). We found that participants continued to improve in both of these metrics as they performed each block of the task. While offline classification metrics for the wrist pose decoder show meaningful rates of misclassification (Fig. 2e), the use of the model in closed-loop for continuous control supported reasonable performance by the metrics presented in Figures 3d and 3e. All participants successfully acquired the target in every trial and the majority of them subjectively reported that the cursor moved in the intended direction >80% of the time (Extended Data Fig. 7c).

**Fig. 3.**
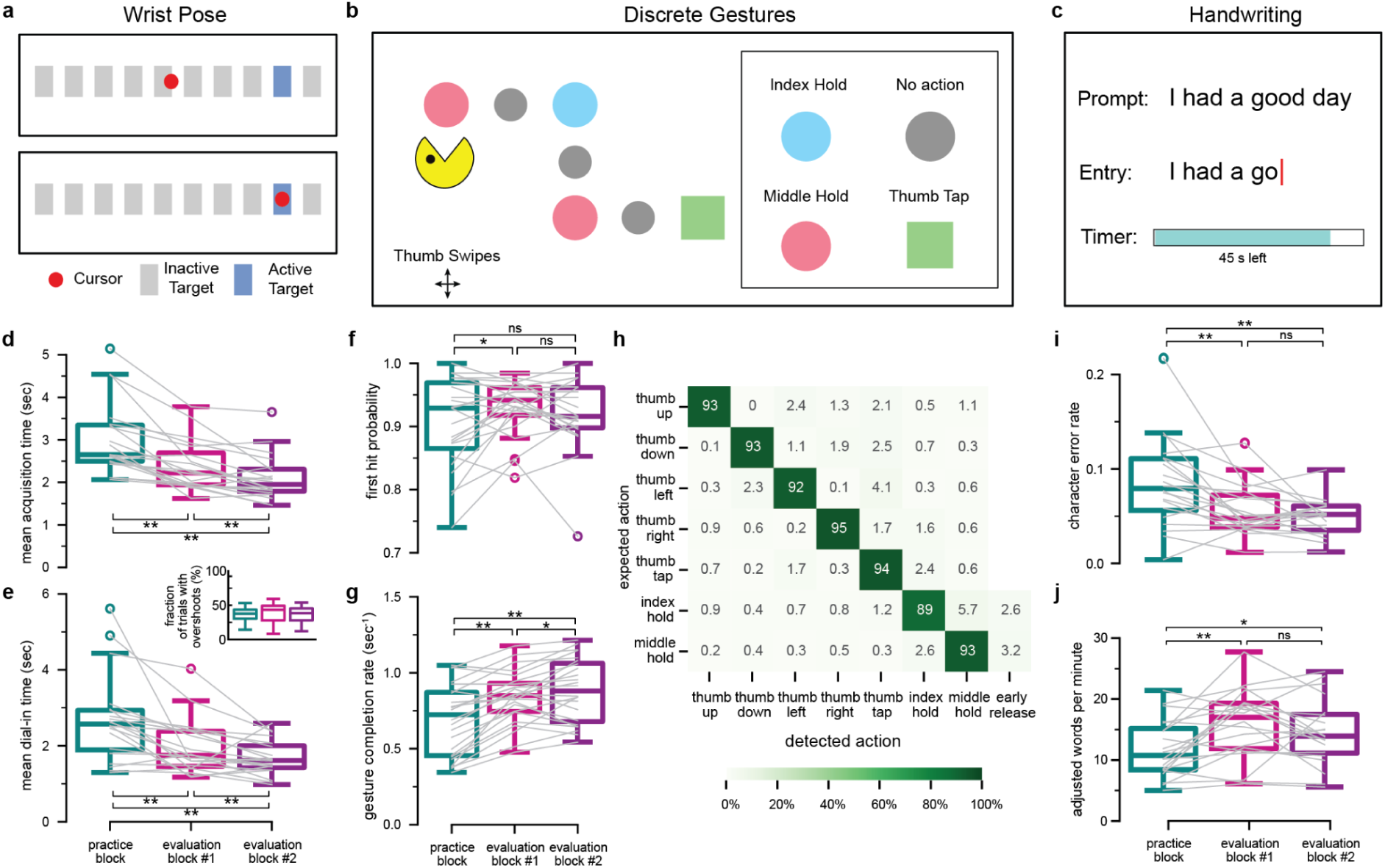
Generic sEMG decoding models enable closed-loop control in diverse interaction tasks. **a**-**c,** Schematics of the three closed-loop interaction tasks demonstrated here. **(a)** A horizontal 1D cursor task in which participants continuously control a cursor (red circle) to acquire a target (blue rectangle) in a row of possible targets (gray rectangles). The participant sees all the possible targets with this trial’s target highlighted; when the target is captured, it disappears and a randomly chosen target among the ones remaining is highlighted to begin the next trial. **(b)** A discrete grid navigation task where participants use discrete thumb swipe gestures to navigate along rectilinear trajectories and perform prompted actions when specified by colored shapes. **(c)** A prompted text entry task in which participants input text via handwriting movements. (See Supplementary Videos 1-4). **d**-**e**, Performance of *N*=20 naive test participants using the wrist-pose decoder to perform a horizontal 1D continuous cursor control task. **(d)** Mean target acquisition time in each block of the task; median in final block is 1.95s. Here and below, boxplots show the distribution over participants and overlaid gray lines show individual participants’ data. **(e)** Mean dial-in time in trials in which the cursor prematurely exited the target before completing the 500ms hold time; median in final block is 1.61s. Inset shows the fraction of trials in which this occurred; median in final block is 38.4%. For both acquisition time and dial-in time, performance in each block significantly improved relative to the previous block, indicating learning effects over time. **f**-**h**, Performance of *N*=24 naive participants on a discrete grid navigation task in which participants were prompted to perform the 9 discrete gestures. **(f)** Fraction of trials in each block in which the first gesture detected by the model was the correct one (“first hit probability”, out of 130 total prompted gestures in each block); median in final block is 0.92. (**g)** Participants’ mean gesture completion rate in each task block; median in final block is 0.88 completions per second. Mean gesture completion rate in each block significantly increased over the previous block, indicating learning effects over time. (**h**) Discrete gesture confusion rates in evaluation blocks, averaged across participants. Confusion rates are expressed as a percent of instances in which the corresponding gesture was expected (across rows). **i**-**j**, Performance of *N*=20 naive test participants on a handwriting task. **(i)** Character error rate (CER) over prompts in each block; median in final two blocks is 0.048 and 0.052. CERs were significantly lower in the evaluation blocks than in the practice block. **(j)** Participants’ aggregate adjusted words per minute (aWPM): ((1-CER) * WPM) over prompts in each block of the handwriting task; median in final two blocks is 17.0 and 13.9 adjusted words per minute. Note that a few outlier prompts with poor performance (e.g. due to accidentally pressing the submit button before completing the prompt) had a strong influence on the aggregate metric for some users; the median CER or aWPM over prompts was respectively lower (0.039 and 0.046 in last two blocks) and higher (19.6 and 17.7) than the aggregate over prompts. For all panels, one and two asterisks respectively indicate *p*<0.05 and *p*<0.005, and “ns” indicates “not significant” (*p*>0.05); one-tailed Wilcoxon signed-rank test.

For discrete gestures, all participants were able to complete the task by navigating with the swipe gestures and performing the activation gestures (thumb tap, index pinch-and-hold, middle pinch-and-hold) when required. Performance on the discrete gesture task was characterized by a measure of how often the first detected gesture following a prompt matched the prompted gesture (Fig. 3f) as well as how long it took to complete each prompted gesture (Fig. 3g). Acquisition rates improved with practice but first hit probabilities did not, suggesting that participants were adapting to the task rather than to the model. The confusion matrix across discrete gestures is shown in Fig. 3h. Note that errors on this task (reflected in both confusions and first hit probabilities) are a combination of model decoding errors as well as “cognitive” errors, where the participant performed the wrong gesture. Index and middle holds were sometimes released too early (i.e. the detected release followed the detected press less than 0.5s later), which are indicated in this confusion matrix as an “early release”. The rate of misclassification seen offline (∼7% mean error over users, Fig. 2f) is comparable to the closed loop confusion rates (∼93% mean accuracy over users, diagonal of Fig. 3h).

Performance of the closed-loop handwriting decoder was evaluated by participants entering prompted phrases and was characterized by character error rate (Fig. 3i) and speed of text entry (Fig. 3j). Improvements from practice to evaluation blocks indicate that participants were able to use the practice trials to discover handwriting movements that were effective for writing accurately with the decoder.

### Representations learned by the discrete gesture model

The neural networks employed as generic sEMG decoding models for this work learn to recognize sEMG patterns based on training with participant data. To confirm our intuitions about how the decoders function, we visualized the representations learned by the intermediate layers of the discrete gestures decoder. The network architecture consisted of a 1D convolutional layer, followed by 3 recurrent LSTM layers (Hochreiter and Schmidhuber 1997) (Fig. 4a), and finally a classification layer (not shown).

**Fig. 4.**
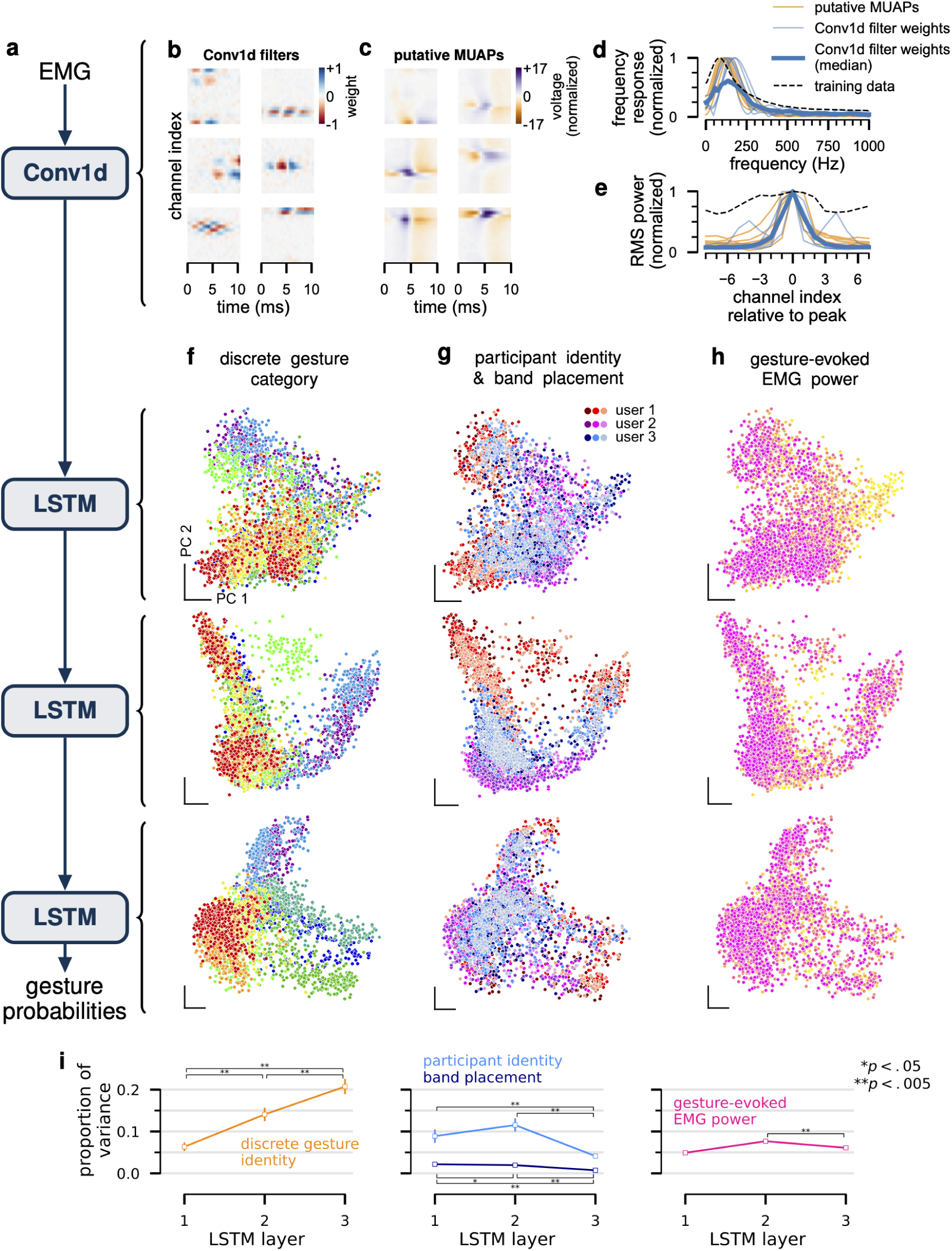
The discrete gesture decoder learns representations that are physiologically grounded and with deeper layers increasingly organized by gesture category and invariant to nuisance variables. **a**, Schematic of network architecture, consisting of a convolutional layer followed by three LSTM layers. Final linear readout and intermediate normalization layers are not shown (Online Methods). **b**, Representative convolutional filter weights from the first layer of the trained model. Filters are typically spatially localized and have a bandpass frequency response. **c**, Example putative MUAPs recorded with the sEMG wristband (after high-pass filtering, following the same preprocessing procedure for the data inputted to the model). **d**, Frequency response, normalized by its peak, of channel with maximum power, for each example convolutional filter from panel b (thin blue lines) and each MUAP in panel c (orange lines). Thicker blue line shows the median over all convolutional filters in the trained network. The full distribution of spectral properties of all the convolutional filters are shown in Extended Data Fig. 8. Dashed black line shows the mean frequency response over channels of sEMG from 10 randomly sampled sessions from the model training set. Note that the sEMG is pre-processed by high-pass filtering (see Online Methods), thus the frequency response cuts off sharply at 40Hz. **e**, Per-channel root mean square power, normalized by its peak, for each example convolutional filter from panel b (thin blue lines) and each MUAP in panel c (orange lines). As in previous panel, thicker blue line shows the median over all convolutional filters in the trained network and dashed black line shows the per-channel normalized RMS power of sEMG over 10 sessions from the model training set. **f**-**h**, PCA projection of LSTM representations of 500ms sEMG snippets aligned with instances of each discrete gesture, from 3 participants held out from the training set, with 3 different band placements. Each row shows the representation of each LSTM layer. Each column shows the same data, colored by: (f) discrete gesture category, (g) participant identity and band placement, or (h) sEMG root mean square power at the time of the gesture. Whereas the discrete gesture category appears to become more tightly clustered at the deeper LSTM layers, nuisance variables like participant identity, band placement, and sEMG power appear to be more clustered at the earlier LSTM layers and less so at the deeper layers. **i**, Proportion of total variance accounted for by each variable, for each layer (mean and 95% confidence intervals estimated from all 50 test participants; see Online Methods). The proportion of variance accounted for by discrete gesture category increases as a function of layer depth whereas that accounted for by the participant identity and band placement nuisance variables decreases. Differences between layers are statistically significant in most cases (*p* < .05, one-tailed paired sample *t*-test) as indicated by asterisks. Error bars (barely visible) show 95% Student-t confidence interval for the mean.

To interpret the convolution layer, we visualized representative filters (Fig. 4b) alongside putative MUAPs (Fig. 4c) detected using the wristband. The filters appear to form a coarse basis set spanning the statistics of MUAPs; specifically, Fig. 4d,e show the general similarity in frequency content and spatial envelope between the putative MUAPs and emergent convolutional filters.

To examine the intermediate LSTM representations, we visualized the changing representational geometry across layers. We analyzed the representations of four properties: gesture category, participant identity, band placement, and gesture-evoked sEMG power (a proxy for behavioral variability over executions of the same gesture). Fig. 4f-h show LSTM hidden unit activity at each layer evoked by snippets of sEMG activity triggered on discrete gesture events, colored by one of the four aforementioned properties. By examining the dominant principal components (PCs), we observed a trend of gesture category becoming more separable deeper in the network as the representations of each gesture become more tightly clustered and less sensitive to nuisance variables (participant identity, band placement, and power). With increasing depth in the network, gesture category accounted for an increasing proportion of the variance in the representation of each layer (Fig. 4i; Online Methods). In summary, the network learns to solve this task by progressively shaping its representation of the EMG to be more and more invariant to nuisance variables.

### Personalizing sEMG handwriting models improves average performance

While generic models allow a neuromotor interface to be used with little to no setup, performance can be improved for a particular individual by personalizing the generic model to data from that participant. We explored personalization for the handwriting task through the fine-tuning of all of the generic model’s parameters using additional supervised data from a set of 40 participants not included in the training data of the generic model. For each participant, we held out a single session that was recorded in each of the three postures used in training (see Online Methods). We then trained personalized models for 300 epochs without early stopping on prompts sampled uniformly at random from all of their remaining sessions (Fig. 5a).

**Fig. 5.**
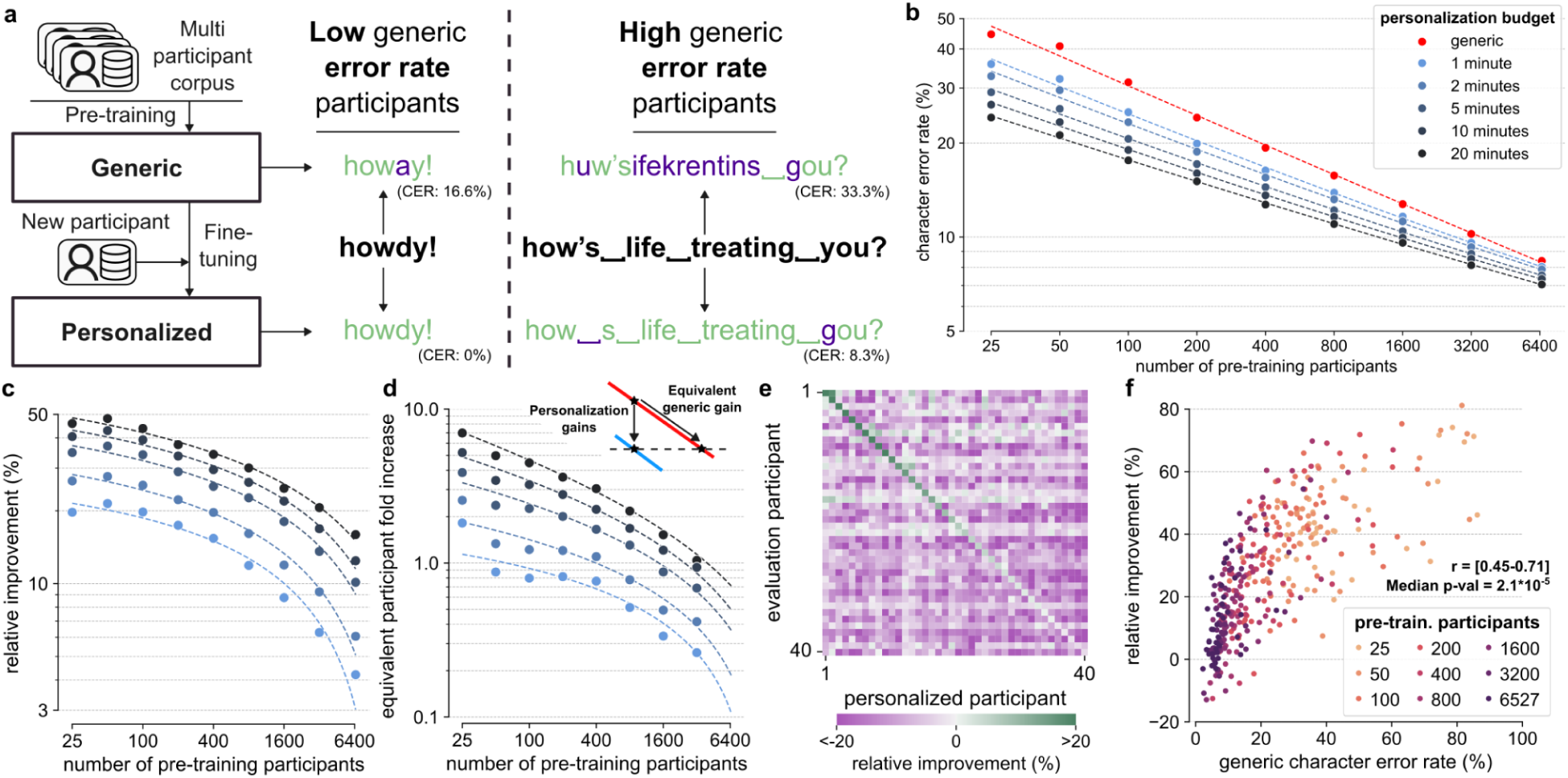
Personalization of sEMG handwriting models improves performance over a generic model. **a**, Schematic of the supervised domain adaptation approach for handwriting personalization. Predictions for example prompts are shown before (above the bold prompt, e.g., “howdy!”) and after personalization (below the bold prompt) for two participants. Prompts are shown in black, correct predictions in green, and incorrect predictions in purple. **b**, The performance of a 60.2M parameter model pre-trained on varying numbers of participants (red). Performance of pre-trained models fine-tuned on varying amounts of personalization data for each test participant (shades of blue). All points represent the population mean of test participants for each setting. The dashed lines show power law fits (Online Methods). **c**, The relative improvement of personalized models increases with increasing duration of fine-tuning data per participant but decreases as the number of pre-training participants increases. The y-axis shows the relative reduction in CER that personalization provides beyond a given generic model, with higher numbers indicating larger improvements. The x-axis and colors are the same as **b**. Dashed lines are the relative improvements calculated from the power law fits shown in **b**. **d**, The relative increase in the amount of pre-training data required to match performance for a given amount of personalization data decreases with larger amounts of pre-training data (Online Methods). The y-axis shows the relative increase in the number of pre-training participants, the x-axis and colors are the same as **b**. Dashed lines are the relative increases calculated from the power law fits shown in **b**. **e**, Personalizing models yields negative average transfer to other participants but can occasionally benefit some participants. Each column shows a single personalized model (on 20 minutes of data), each row a single test participant’s data, and the colors the relative reduction in CER of that model from generic model performance on that participant’s data (positive, green, is an improvement). Diagonals show improvement due to personalization on the same participant. We use the 60.2M parameter model pre-trained on 6527 participants as base for personalization. Columns and rows are sorted by participants most improved by personalization (diagonal). **f**, Across the population of test participants, those with worse performance on the pre-trained model tend to exhibit larger relative improvement, and the best performing users occasionally exhibit regressions. We show the relative reduction in CER (higher is better) of the 60.2M parameter model fine-tuned on 20 minutes of data without early stopping. Results are shown for all numbers of pre-training participants. Each dot represents one participant (participants are repeated across pre-trained models). The x-axis shows generic model performance for each model. We show the range of Spearman correlation coefficients for each fit and the median p-value (maximum p=0.0035).

Fine-tuning generic models improved their average CER for all amounts of additional data and for all numbers of pre-training participants (Fig. 5b). In all cases, more personalization data led to further reductions in population average CER. However across all generic models, as the generic model was pre-trained with data from more participants the absolute and relative improvement in CER from personalization diminished (Fig. 5c), indicating that there are diminishing returns to personalizing already performant generic models.

Personalizing models can thus be seen as an alternative to expanding the generic corpus size in order to decrease a model’s CER on the target participant (Fig. 5d). For example, for the model pre-trained on the smallest corpus of 25 participants (or 1,900 minutes), personalization with 20 minutes of data from the target participant was equivalent to training a generic model with 14,000 minutes of additional data from other participants—effectively adding 7x as much data as in the original pre-training corpus. For the model trained on 3,200 participants (or 240,000 minutes), 10 minutes of personalization led to a model as performant as a generic model pre-trained on double the number of pre-training participants. Thus as more data from other participants is added, the effective enhancement of the generic training corpus achieved through personalization diminishes. Conversely, fixed enhancements of the generic training corpus correspond to a diminishing amount of personalization data—adding 14,000 minutes of data is equivalent to 20 minutes of personalization for the 25 pre-training participant model and only about 1 minute for the 200 pre-training participant model.

While personalization improved performance on the target participant, model performance improvements from personalization did not transfer across participants. For the most performant generic model trained (6,527 participants, 60.2M parameters), we found that personalizing on one participant and evaluating on another participant generally had a negative impact on performance when compared to the generic model performance (Fig. 5e). Personalization on the same participant improved performance in 88% of participants and led to a relative improvement of 8.35% ± 2.36% (median ± s.e.m. over participants), while personalization on a different participant (data from one participant used to personalize a model for another participant) only improved performance on 7% of such participant pairs and led to an average relative decrease of 8.86% ± 0.53% (median ± s.e.m., taken across each evaluation participant after averaging across personalized models, Online Methods).

Personalization disproportionately improved the performance of poorly performing participants across all generic models (Fig 5f). For example, for generic models pre-trained with 6,527 participants, personalization provided larger relative gains for participants with higher generic model CER (Fig. 5f, inset) and more moderate gains or occasional regressions for those with already low CERs. (In Extended Data Figure 9, we show that these regressions can be mitigated with early stopping during fine-tuning, albeit at the cost of increased data required for validation.)

The degree to which models can be personalized depends on the ability to collect fine-tuning data in a given deployment environment, the need for the performance gains from personalization, and the ability to collect pre-training data. Overall, these results define clear trends and tradeoffs for personalization that enable rational design of data collection and modeling strategies for neuromotor interfaces.

## Discussion

We introduced an easily donned/doffed wrist-based neuromotor interface capable of enabling a diverse range of computer interactions for novel users. We began by developing a robust and scalable data collection framework that allowed us to collect large training corpora across diverse participants (Fig. 1). We then used contemporary supervised deep learning to produce generic sEMG models that overcame issues observed in single-participant models that have long stymied generalization in brain computer interfaces (Fig. 2). The resulting sEMG decoders enabled continuous control, discrete input, and text entry in closed-loop evaluations without any need for session- or participant-specific data or calibration (Fig. 3). A dissection of intermediate representations in the discrete gesture neural network decoder highlighted its ability to disentangle nuisance parameters related to band placement and behavioral style (Fig. 4). Finally we demonstrated improvements in handwriting decoder that increase with ever more personalization data, suggesting that models can continue to improve through model refinement or scaling data collection (Fig. 5). Together this work defines a framework for building generic interaction models using non-invasive biological signals.

### Related work in HCI and BCI

The work presented here sits at the nexus between HCI and BCI, building a bridge between these fields. The HCI community has placed significant emphasis on advancing gestural input for various technology applications by deploying machine-learning-backed solutions for differing sensing modalities such as computer vision (Kinect, Meta Quest, etc.) inertial measurement units (Wen, Ramos Rojas, and Dey 2016), sEMG (Mendez et al. 2017; Saponas et al. 2008), bio-acoustic signals (Laput, Xiao, and Harrison 2016), electrical impedance tomography (Zhang and Harrison 2015), electromagnetic signals (Laput et al. 2015), and ultrasonic beamforming (Iravantchi, Goel, and Harrison 2019). The most direct antecedent of the work presented here is the now discontinued sEMG Myo armband (worn on forearm) by Thalmic Labs, as used for gesture detection by e.g., (Mendez et al. 2017), who built single-user models to classify more intuitively separated gestures such as wrist extension, open hand, etc. We demonstrated our sEMG-RD’s advantages over existing HCI work in three main areas: self-contained sensing suitable for mobile environments, generalizable and performant algorithms trained on thousands of participants for cross-user generalization, and an approach that encompasses a broader spectrum of recognition, including handwriting, gestures, and poses to input text and discrete and continuous controls.

Our non-invasive sEMG work has intimate connections to BCI. EEG-based BCI systems lag significantly behind other modalities in terms of performance for a variety of reasons, primarily relating to issues with signal quality and secondarily interpretation, but also issues such as a lack of unification around hardware or software (Jayaram and Barachant 2018). As a result, modeling efforts have been focused on small models and relatively small datasets (e.g. <50 users, (Lawhern et al. 2018)). Beyond EEG-based BCI, the use of EEG to analyze sleep has more recently deployed neural networks on tens of thousands of participants (Perslev et al. 2021) and EEG-based BCIs (notably, spellers) can still achieve impressive bitrates of 100-300 bits per minute using variations of the steady-state visually evoked potential paradigm (Maslova et al. 2023) (our handwriting decoder showed a per-minute bit rate of 528 bits per minute). However, these EEG systems—while highly optimized for speed—require individualized setups and highly intrusive display elements.

In the field of intracortical BCI, approaches have so far been limited to single participant studies and hindered by nonstationarities in recordings and over sessions (Degenhart et al. 2020; Brandman et al. 2018; Sussillo et al. 2016; Gilja et al. 2012). While the field of intracortical BCI is transitioning to neural network decoders (Sussillo et al. 2016, 2012; Willett et al. 2023; Metzger et al. 2023), it remains focused on solving these calibration issues, which result largely as a function of limited data. Given that sEMG signals derive from the summed activity of motor unit firing, it is possible that sEMG decoding methods can guide methods developed for intracortical BCI systems, despite the fact that sEMG is completely noninvasive with more direct correspondence between recorded signals and motor behavior. The large-scale approaches demonstrated here may provide direction to the larger BCI field as it scales up beyond single-user studies, e.g. (Musk and Neuralink 2019).

As a point of comparison, (Willett et al. 2021) describe an invasive BCI for decoding imagined handwriting that achieves mean text entry speeds of 18 words per minute and error rates of 5.9%. This is the fastest character-based BCI for text entry in the literature. The results reported here for our non-invasive method are generally similar, with a median speed over users of 17.7 words per minute (aggregate over prompts, median over prompts was 20.9 due to outliers) and median CER over users of 4.8% (aggregate over prompts, median over prompts was 3.9%). However, we emphasize that the (Willett et al. 2021)) system requires surgical implantation of electrodes and 5 days of practice, whereas our focus is on non-clinical populations and only requires non-invasive electrodes and minutes of practice. Our results are not yet as performant as typing on a mobile keyboard, for which (Palin et al. 2019) measure speeds for 37,000 users of an online typing test tool with users’ own devices that could include autocorrect and other assistive technologies. They find an average typing speed of 36.2 words per minute and an uncorrected error rate (percentage of errors remaining after user editing) of 2.3%. Finally, the fastest text entry BCI is described by (Willett et al. 2023) and operates at the word-level on attempted speech from a subject with amyotrophic lateral sclerosis via invasive intracortical recordings. This system was able to achieve speeds of 62 words per minute with a 23.8% word error rate on a vocabulary of 125,000 words. While our current noninvasive sEMG-based handwriting interface operates at the character-level, word-level models achieving faster writing speeds could be feasible though would require users to learn novel schemes.

### Datasets and model fine-tuning

Our results suggest that collecting large corpora of diverse data provides a means for constructing generalized sEMG models that are performant across a diverse population. We recapitulated the limitations of single-participant models (Fig. 2) and showed that generalization can be achieved using sufficiently large, diverse datasets and high-capacity deep learning models (Fig. 3d-k). In particular, we observed power law scaling of model performance with the number of participants recorded (Fig. 3a-c). These power laws mirror emerging work in natural language processing and computer vision that highlights power law scaling of network performance with the amount of data recorded (Hoffmann et al. 2022; Kaplan et al. 2020).

The fine-tuning of models on data from a specific individual further improved system performance for that individual (Fig. 5). While generic model performance may suffice for most users, this form of personalization can be applied selectively to users for whom the generic model works insufficiently well due to differences in anatomy, physiology, or behavior. Personalization has shown benefits to accuracy for automatic speech recognition in language models (McGraw et al. 2016) and acoustic models (Mdhaffar, Tommasi, and Estève 2021) as well as speech enhancement (Eskimez et al. 2022). (Pillutla et al. 2022) have shown that federated learning is able to partially personalize models in the field in a privacy preserving manner that does not send user data to a centralized repository, so it is possible that the user benefits of such approaches could be achieved purely on a user’s own device.

### Future directions

This non-invasive, wrist-based interface allows direct intentional motor signal detection from the muscle; thereby, the sEMG decoder opens new directions in novel and accessible computer interactions. For instance such a decoder could be used to directly detect an intended gesture’s force, which is generally unobservable with existing camera or joystick controls. Moreover the sensitivity of sEMG to detect signals as subtle as putative individual MUAPs (Fig. 1b) enables the creation of extremely low-effort controls—an important innovation with potential impact for people with a diverse range of motor abilities or ergonomic requirements. Explorations of interactions in neuromotor signal space—as opposed to gesture space—may enable entirely novel forms of control, for example by exploring the limits of novel muscle synergies or interaction schemes that directly depend on individual motor unit recruitment or firing rate control.

Scientifically, a viable research platform, such as sEMG-RD, could enable studying, for example, the effects of neurofeedback on motor unit activity for novel human-machine interactions (Formento, Botros, and Carmena 2021; Bräcklein et al. 2022, 2021), decoding of motor function (Stachaczyk et al. 2020; Tanzarella et al. 2023), or investigating the underpinning mechanisms of motor unit control (Marshall et al. 2022; Hug et al. 2023).

Finally, in the clinic, the ability to design interactions that require only muscular activity rather than performance of a specific movement or interaction with a keyboard or joystick could enable viable interaction schemes for those with reduced mobility, muscle weakness, or missing effectors entirely (Yamagami et al. 2023). All of these novel interactions will be facilitated by continued improvements in the channel count and sensitivity of sEMG devices, and potentially other signals recorded at the wrist.

## Supporting information

Supplemental Video 1 Wrist Rotated

Supplemental Video 2 Discrete Rotated

Supplemental Video 3 HW Rotated

## Contributions

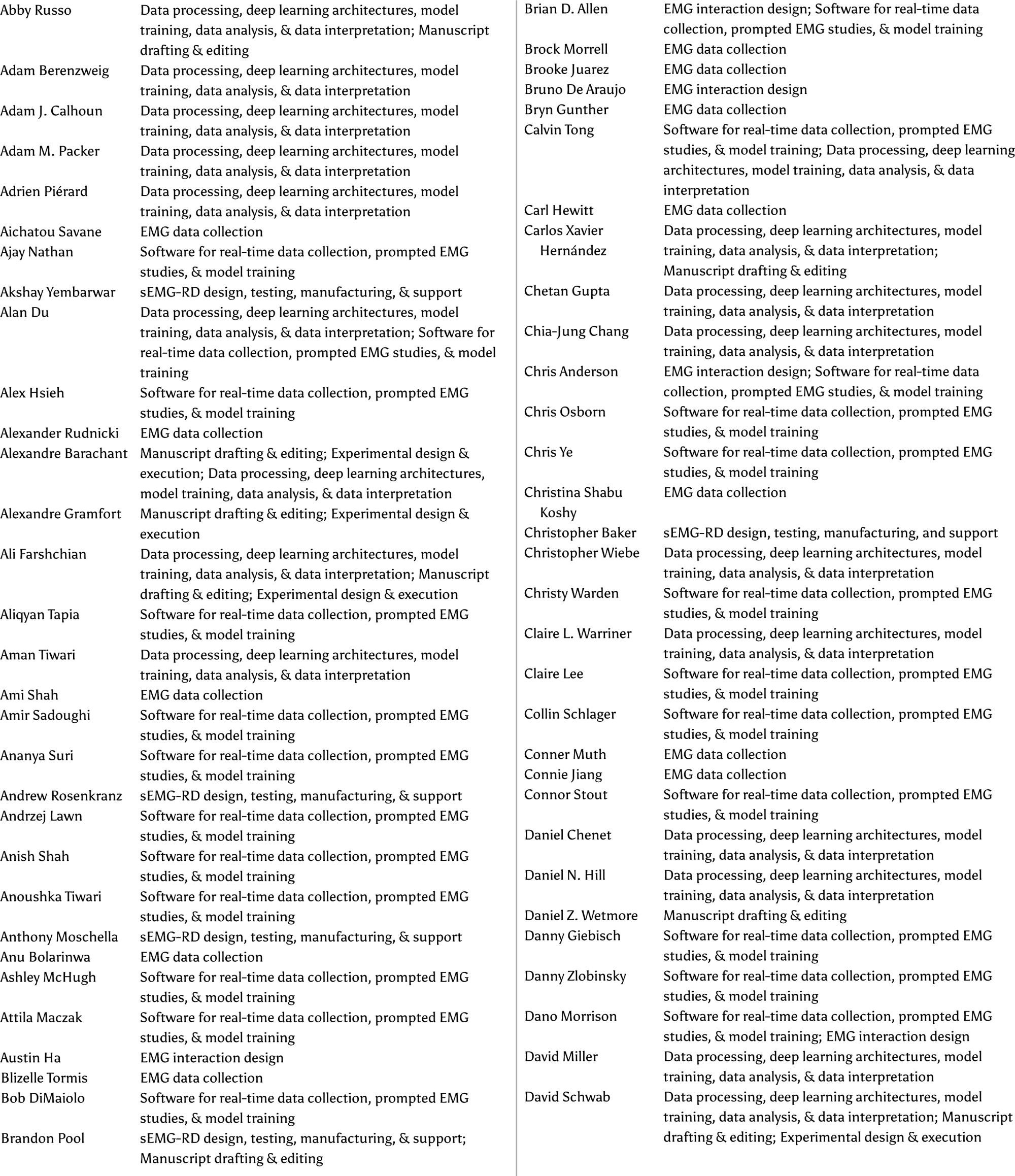
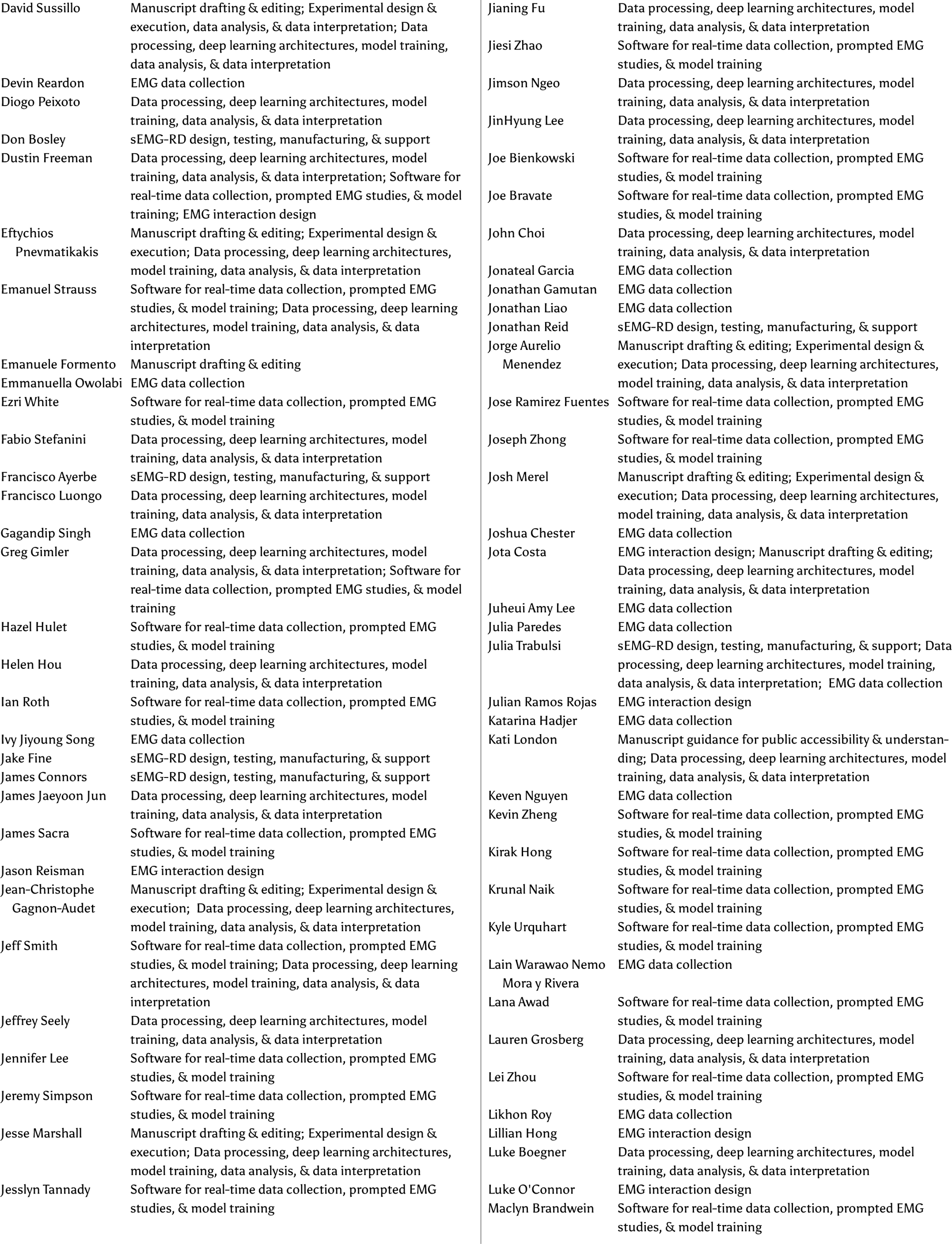
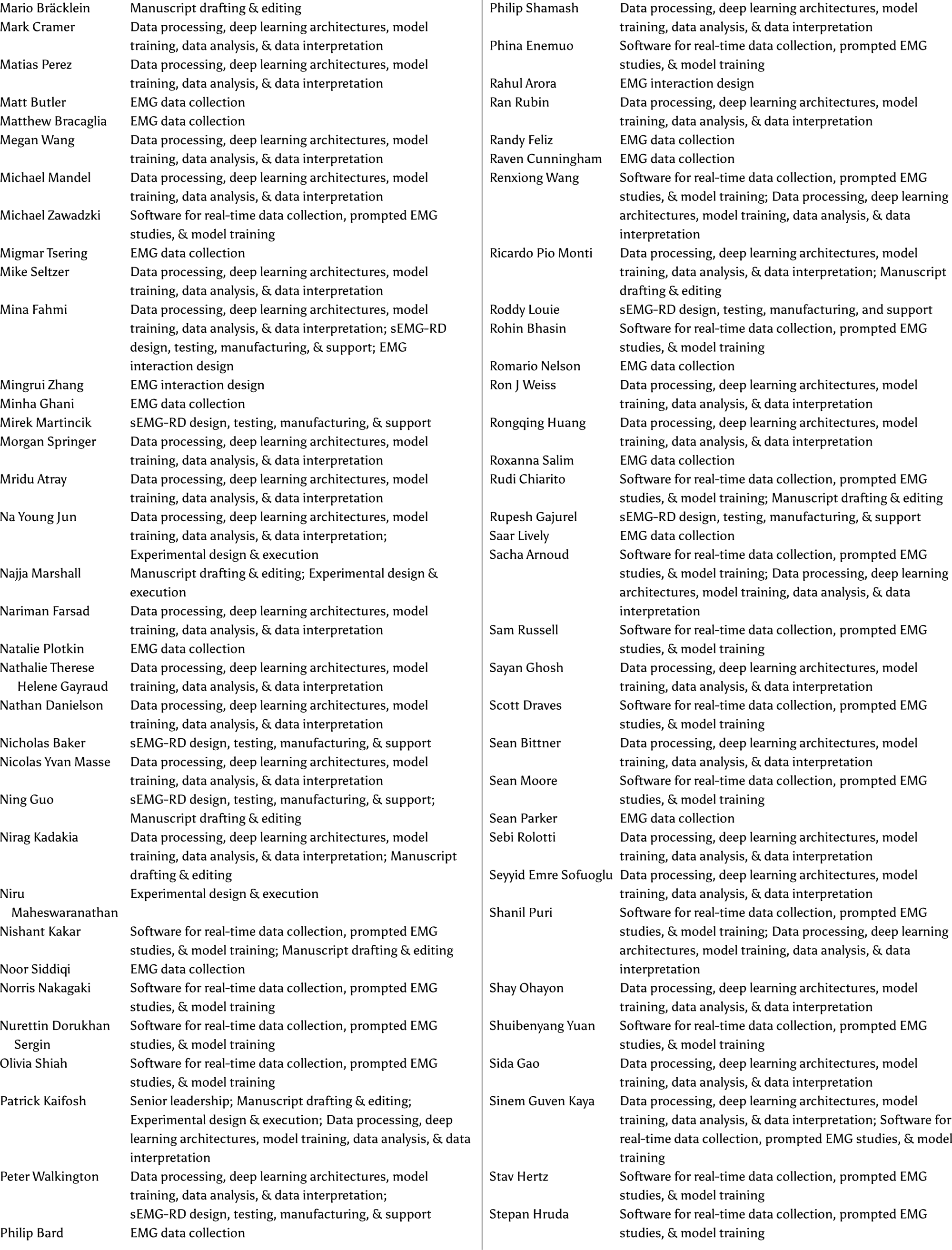
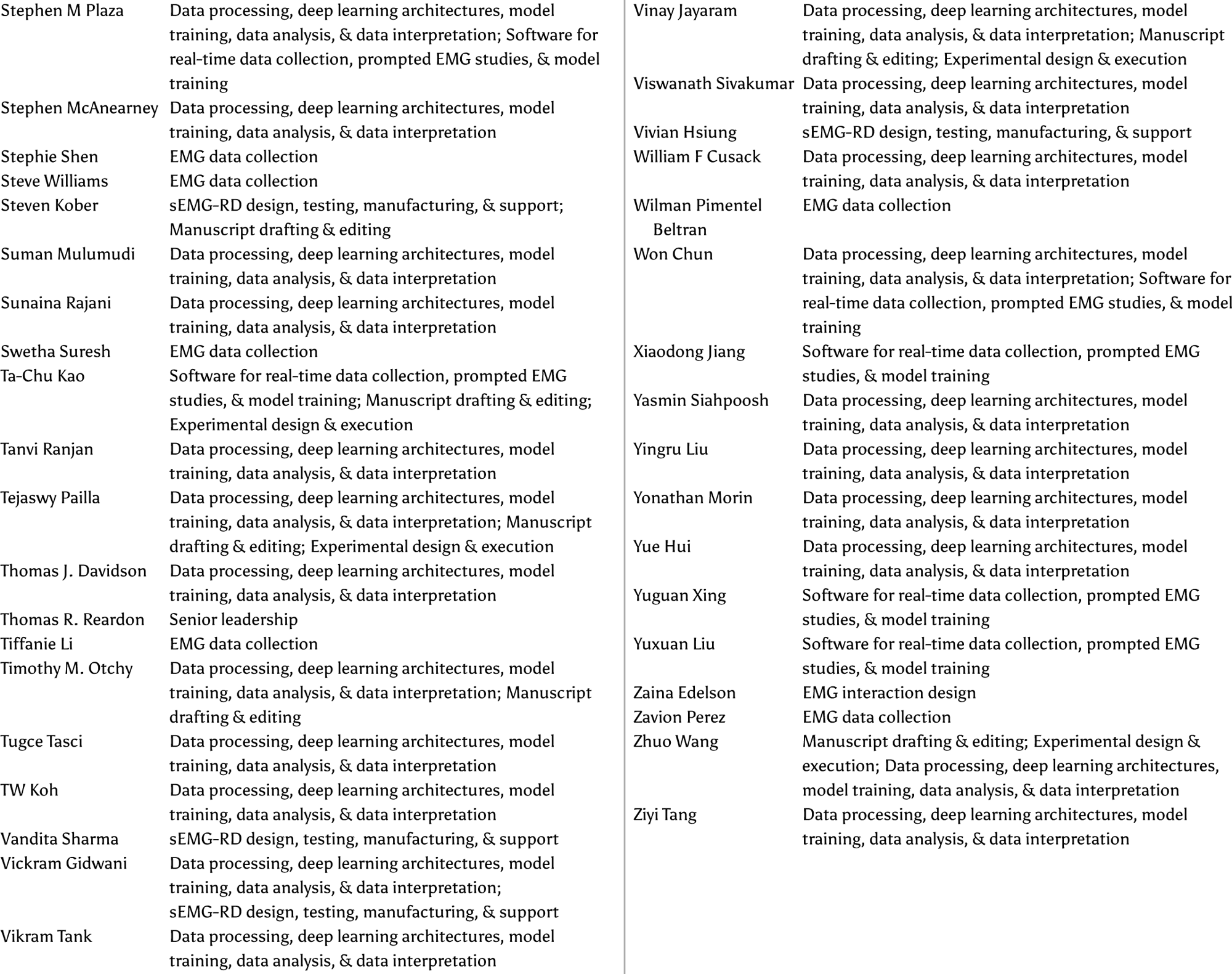

## Acknowledgements

We thank Shaul Druckmann, Dario Farina, Adrian Haith, John Krakauer, Amy Orsborn, Chethan Pandarinath, Liam Paninski, Krishna Shenoy, and Daniel Wolpert for technical and scientific advice and feedback. We thank Dario Farina, Deren Barsakcioglu, and Pedro Rente Vicente of Imperial College London for assistance with the wrist MRI scans.

## Online Methods

### Hardware

#### sEMG-RD Device

The surface electromyography devices consisted of two primary subcomponents: a digital capsule and an analog wristband (Extended Data Fig. 1). The digital capsule comprised the battery, antenna for Bluetooth communication, and a printed circuit board that contained a microcontroller, an analog-to-digital converter, and an inertial measurement unit. The analog wristband comprised discrete links that each housed a multi-layer rigid printed circuit board that contained the low-noise analog front-end circuits and gold-plated electrodes. The analog front end applied 20 Hz high-pass and 850 Hz low-pass filters to the data.

These printed circuit boards were inserted into Nylon 12 PA 3D printed housings and then strung together with a multi-layer flexible printed circuit board along with a strain-relieving fabric. An elastic nylon cord was routed continuously between the links and was tied together at the wristband gap to form a clasp and tensioning mechanism. Finally the digital capsule was connected to the analog wristband through a connector on the flexible printed circuit board and fastened together with screws for mechanical stability. The device underwent a biocompatibility testing process to ensure its safety.

### Data collection and processing

#### MRI scan

In order to visualize the position of the sEMG-RD’s electrodes relative to wrist anatomy we collected a high-resolution anatomical MRI scan (Siemens Magnetom Verio 3T) from a consenting participant’s right forearm. We collected axial scans along the forearm, beginning from just distal to the wrist and ending just distal to the elbow. The scan was collected pursuant to an IRB governed study protocol conducted by Imperial College London.

#### Participant Experience

Study recruitment and participant onboarding followed IRB-approved protocol(s), which included providing participants with information about the study protocol and asking them to review and sign an IRB-reviewed consent form prior to beginning the study. Participants were provided with the opportunity to ask questions prior to their participation and were able to discontinue their participation at any time. On-site research administrators monitored participants during the study protocol(s) to ensure participant well-being. Participants were financially compensated for their time participating in the study.

#### Collection at scale

Participants visited data collection and lab facilities to perform the study protocols. On a given day, there were up to 300 participants who partook in a study. Once a participant was in the facility, measurements of the wrist and hand were taken, including forearm circumference and wrist circumference. Next, the participant was fitted with an sEMG-RD band around the wrist to collect sEMG data. Instructions were provided for gestures and movements prompted in the subsequent study session expected to last around 3 hours (including rests and briefing).

All responses and information provided during the study were collected and stored using de-identification technique(s) in a secure database.

#### Prompted study design

All of our tasks were framed as supervised machine learning problems. Insofar as the behaviors we prompted involved overt movements, we could hypothetically leverage instrumentation to obtain ground-truth labels, which would complement the prompted labels. We experimented with various modalities of ground truth labels; however, instrumentation also imposed constraints on what could be explored (dedicated sensors need to be built for each individual modeling task) and limited ecological validity of the signal (physical sensors can restrict movement range, poses, conditions, etc). Therefore we relied on experimental design and prompting to obtain approximate ground truth for our data, rather than direct instrumentation. We acknowledge the potential tradeoff between quality of labels and scalability of the data collection process.

Training and evaluation protocols were implemented in a custom, internal software framework that leveraged the capabilities of Lab.js, an established open-source, web-based study builder (Henninger et al. 2022). This framework orchestrated both the presentation of task-specific prompter applications and the recording of annotations from these applications. The framework was developed using TypeScript and the task-specific prompters were built upon the React framework.

#### Real-time data collection system

Data collection for our studies was performed using an internal framework for real-time data processing that supports data collection, offline model training and benchmarking and online evaluation. At its core, the framework offers an engine for defining and scheduling a data processing graph. On the periphery, it provides well-defined APIs for real-time performance monitoring and interaction with consumer applications (e.g. prompting software, applications for stream visualization).

For data collection, our internal platform served as the host for recording real-time signals and annotations to a standardized data format (i.e. HDF5). For offline model training and benchmarking, our internal platform provides an API for batch processing of data corpora. This helps to generate featurized data from the recorded raw-signals and apply model inference for offline evaluation. To ensure online and offline parity, the internal platform also supports running the same sequence of processing steps on real-time signals for online evaluation.

#### Training data corpora

##### Wrist pose corpus

The wrist pose training corpus included sEMG recordings from 389 participants (3 recording sessions per participant). In each session, a participant repeatedly held 5 wrist poses: wrist flexion, wrist extension, wrist ulnar deviation, wrist radial deviation and wrist neutral. Each pose was performed 8 times, with the order of the poses randomized. The hold times for each pose was between 2 and 5 seconds drawn uniformly at random. After holding each pose, the participant rested between 0.5 and 1 seconds (drawn uniformly). The participants performed these movements in the “laser-pointer” posture with a loose fist in front of the body, thumb at top and elbow at approximately 90 degrees.

##### Discrete gesture corpus

The discrete gesture training corpus was composed of data from 4800 participants. As noted in the main text, there were nine prompted gestures: index and middle finger presses and releases, thumb tap, and thumb left/right/up/down swipes. Each session consisted of stages in which combinations of gestures were prompted at specific times. These combinations usually included the full set of trained gestures, but in some stages were restricted to specific subsets (e.g. pinches only, thumb swipes only). During data collection for these stages, participants were asked to hold their hand and arm in one of a range of postures or to translate/rotate their arms while completing gestures. In ∼10% of stages, instead of prompting specific timing, participants were asked to complete sequences of 3-5 gestures at their own pace. About ⅓ of the training corpus was composed of a range of null data in which participants were either asked to generate specifically-timed null gestures (e.g. snaps, flicks) or to engage in more loosely-prompted longer form null behaviors (e.g. typing on a keyboard).

##### Handwriting corpus

The handwriting recognition corpus comprised sEMG recordings from a total of 6627 participants. The data was collected in short blocks, during which participants were prompted to write a series of randomly selected items, including letters, numbers, words, random alphanumeric strings, or phrases. The set of phrases used for sampling was sampled (after filtering to remove offensive words and phrases) from a dump of Simple English Wikipedia in June of 2017, the Google Schema-guided Dialogue Dataset (Rastogi et al., 2020), and the Reddit corpus from ConvoKit (Chang et al., 2020). Each participant contributed varying amounts of data, but approximately one hour and 15 minutes each on average. Each block was performed in one of three randomly chosen conditions: seated writing on a surface, seated writing on their leg as the surface, or standing writing on their leg. Note that we did not have ground truth information about the fidelity with which participants wrote these prompts, but for a subset of participants, handwriting was performed with a Sensel (Sunnyvale, CA) touch surface device. Visual examinations of a subset of the Sensel recordings suggested that approximately 98% of prompted characters were executed successfully.

### sEMG processing and analysis

#### Putative motor unit action potential extraction

Fig. 1b shows MUAPs evoked by subtle contractions of the thumb and pinky extensors in one participant. For each digit, the participant selected the sEMG channel with maximum variance during sustained contractions based on visual inspection of the raw signals. Subsequently, the participant was prompted to alternate between resting and performing sustained contractions of the chosen digit for three repetitions while receiving visual feedback about the raw sEMG signal on the selected channel. Each rest and movement prompt was 10 seconds long with 1 second inter-prompt intervals. The participant used the visual feedback on the selected channel to titrate the amount of generated force to recruit as few MUs as possible with each contraction.

MUAPs were extracted offline for each digit using a simple spike detection algorithm. The sEMG traces were first pre-processed by filtering with a 2nd order Savitzky-Golay Differentiator filter. The filtered sEMG was rectified to improve the alignment of detected MUAPs, averaged over channels, then smoothed with a 2.5ms Gaussian filter to obtain a 1D sEMG envelope. Spikes were detected via peak finding on the sEMG envelope using `scipy.signal.find_peaks` with prominence=0.5 (Virtanen et al. 2020). MUAPs were extracted using a 20ms long window across all sEMG channels, centered on each peak. The waveforms shown in Fig. 1b were obtained from the selected channel for thumb (12; blue) and pinky (14; pink) extension using all MUAPs detected during the second prompted movement period; no attempt was made to cluster MUAPs into different units. For visualization, the opacity of the each trace was scaled as 1 / (1 + |a_i_ + median(*a*)|), where a_i_ is the peak-to-peak amplitude of the *i*^th^ MUAP and *a* are the amplitudes of all detected MUAPs for each contraction.

#### Multivariate power frequency (MPF) features

The wrist pose and handwriting tasks used custom features extracted from the raw sEMG; we refer to this feature set as multivariate power frequency (MPF) features. To obtain these features, we first applied a 40 Hz high-pass filter. We then extracted the squared magnitude of the cross spectral density with a rolling window of 160 sEMG samples and a stride of 40 samples. The cross spectral density was chosen to preserve cross-channel relationships in the spectral domain. The magnitude of cross spectral density was estimated with a discrete Fourier transform (64 sample, stride of 10) and binned into 6 frequency bins (0-62.5, 62.5-125, 125-250, 250-375, 375-687.5, 687.5-1000Hz). This produced a set of 6 symmetric and positive definite 16×16 square matrices that update every 40 samples, for an output frequency of 50Hz. Building on robust results in the EEG space for this class of features, each of these matrices was then projected into their respective Riemannian tangent space (Barachant et al. 2012). Finally, the first three off-diagonals were preserved and half-vectorized for each matrix, and then concatenated across the 6 frequency bins, producing a single 384 (6×3×16) dimensional vector for each 80ms window. An implementation for both the cross spectral density estimation and tangent space mapping can be found in the pyRiemann python toolbox (Barachant et al. 2023).

#### Discrete gesture time alignment

Since all discrete gesture data collection was performed by prompting participants, we only had access to approximate timing of the gesture execution (i.e. the time at which the participant was prompted to perform the gesture). However, training sEMG decoding models to infer when the participant performs a gesture required more precise alignment of labels with the signal in order to be effective. While a task like handwriting used an alignment free loss (i.e. Connectionist temporal classification, CTC) and would be applicable in this task as well, forced-alignment allowed us to gain much finer control over the latency of the detections produced by our models, which was critical for practical use of discrete gesture as control inputs.

When gestures were well isolated, i.e. when the inter-gesture interval was greater than the uncertainty of the timing, existing solutions from the literature could be readily deployed on sEMG data, leading to robust inference of gesture timing (Williams et al. 2020). However, realistic data collection involved rapid sequence of gestures in close succession, which made identification of timing of individual gestures a challenging problem and required a dedicated solution. Therefore we developed an approach to infer precise timing of the gestures.

Our approach was to infer timing of all gestures in a sequence, defined as a series of consecutive gestures where uncertainty bounds overlap. We did this by searching for the sequence of gesture timings that best explained the observed data according to a generative model of our MPF features.

First, for the purposes specifically of this timing adjustment stage, we defined the generative model for a set of K gesture instances, as the sum of gesture-specific templates centered at corresponding event times *t_k_*, with additive noise:

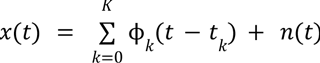

Where *x(t)* is our features over time, ϕ*_k_* (*t*) is a prototypical spatio-temporal waveform for gesture of index *k* (i.e. the gesture template for the class of gesture corresponding to event k), and *n*(*t*), a noise term. We note that this generative model is only valid for ballistic gesture execution and power-based features. We also note that templates are shared across execution of the same gesture type.

We define the *generative inference* as the joint optimization of gesture templates and times at which each gesture occurred. We solved this via an expectation-maximization algorithm (Dempster, Laird, and Rubin 1977): we first estimated the templates based on prompted times, then inferred timestamps of the gesture sequence, and repeated with new inferred event times until convergence, (i.e. when the timestamp updates across iterations of the EM algorithm were smaller than a tolerance value).

Templates were estimated by an EMG analog of the regression-based estimator of the event-related potential (rERP; (Smith and Kutas 2015), in order to disentangle overlapping contributions of gesture when performed in fast sequence. We note here that the gesture templates ϕ*_k_* (*t*) are specific to each participant and band placement. Timings were obtained by the following optimization problem:

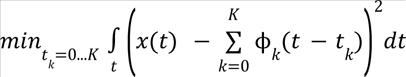

We optimized this numerically through a beam search algorithm, subject to additional *ad hoc* constraints that bounded how far the adjusted times could deviate from the prompted times based on priors from the data collection protocol.

Direct application of the above procedure produced timestamps that were referenced to the session template, and there was an indeterminacy as to the timing offset within the gesture, which can vary due to initial conditions. To better standardize alignment of template timing across individuals, we performed a global recentering step at the end of timestamp estimation. Specifically, we found the time of maximal correlation between the session template (i.e. for a particular participant) and a global template (grand average of all templates across participants).

#### Additional details for single-participant models

##### Training details

To train the single-participant models for the discrete gesture classification task we selected 100 participants that had completed at least 5 sessions of data collection and selected 5 of those sessions. Then we randomly picked 4 of these sessions for training and the remaining held-out session for testing. From these 4 sessions we randomly created nested subsets of 2, 3, and all 4 sessions to train 3 different models for each participant. Given the limited amount of training data per model, we use the MPF feature and a small neural network as described below.

##### Architecture

The single-participant discrete gestures model took as input the MPF features. The network architecture consisted of (a) one fully-connected (FC) layer with Leaky ReLU followed by (b) cascaded time-depth separable (TDS) blocks (Hannun et al. 2019) across time scales and (c) three more FC layers to produce a logit value for each of the 9 discrete gestures to be predicted. For (b), we use two TDS blocks per time-scale: at each scale *s*, an AveragePool layer with kernel size 2^*s* is applied to the output of (a) and fed to a TDS block with dilation 2^*s*. The output is then added to the output of scale *s*-1 (if it exists) and passes through another TDS block with dilation 2^*s* as the output of scale *s* to be used by the next scale *s*+1 (if it exists) or subsequent layers. We used 6 scales (*s*=0,…,5), and the feature dimension was set to 256 for all TDS blocks and all but the very last FC layer.

##### Optimization

We used the standard Adam optimizer with the following learning rate schedule: the learning rate increases linearly from 0 to 1e-3 over a 5-epoch warm-up phase, then undergoes a one-time decay to 5e-4 after epoch 25, and remains constant thereafter. Each model was trained for 300 epochs to avoid under- or over-fitting for single user models based on our experience. Training was done using a binary cross-entropy loss.

#### Related deep learning architectures and approaches

The three HCI tasks described here, continuous pose prediction, discrete action recognition, and the transcription of handwriting into characters, represent related but distinct time series modeling and recognition tasks. Machine learning, and specifically deep learning, approaches have become extremely popular solutions to these problems, including convolutional models (e.g., Karpathy et al., 2014), recurrent neural networks (e.g., Graves et al., 2013), and streaming transformers (e.g., Gulati et al., 2020).

As an example of the similarity between our tasks and established machine learning problems, consider the relationship between handwriting recognition from sEMG and automatic speech recognition (ASR) from audio waveforms. Both tasks map continuous waveform signals (with dimensionality equal to the number of microphones or sEMG channels) at a fixed sample rate, to a sequence of tokens (phonemes or words for ASR, characters for our sEMG-RD). Components of our modeling pipeline have analogs in ASR, including: feature extraction, data augmentation, model architecture, loss function, decoding, and language modeling. As noted below, each of these modeling pipeline components required substantive domain-specific modification for sEMG models. For feature extraction, ASR typically uses log mel filterbanks; we used our analogous multivariate power frequency features (Online Methods). For data augmentation, we used the ASR technique of SpecAugment (Park et al., 2019), which applies time- and frequency-aligned masks to these spectral features during training. A popular model architecture for ASR is the Conformer (Gulati et al., 2020), which provides the advantages of attention-based processing in a form that is compatible with causal time series modeling. We find that this method worked well for sEMG-based handwriting recognition as well. A popular loss function for ASR is CTC (Graves et al., 2006), which allows neural networks to be trained from pairs of waveforms and their textual transcriptions, without the need for a precise temporal alignment. As we similarly had pairs of sEMG recordings and transcriptions without precise temporal alignment, we also utilized CTC to train our models. When decoding models at test time, ASR employs a beam search (Ney et al., 1989) to approximate the full forward-backward algorithm lattice (Rabiner, 1989) while still incorporating predictions from a language model, biasing decoding towards more likely character and word sequences. We similarly integrated beam decoding with language modeling into our decoders (e.g., Pratap et al., 2020), although in the experiments we used “greedy” CTC decoding for the sake of simplicity.

#### Wrist pose

To train wrist pose classification models, we focused on three wrist poses, which corresponded to a subset of the prompted data: wrist flexion, wrist extension, and wrist neutral. We excluded wrist ulnar deviation, wrist radial deviation, and rest, by masking out time points corresponding to those poses in the loss function (cross-entropy loss). We consistently held out a fixed set of 20 participants for validation and 39 participants for testing, and varied the number of training participants from 20 to 320. We used a batch size of 1024, with each sample in the batch consisting of 3 (contiguous) seconds of recordings. When evaluating the model’s offline performance, we took the most common prediction in the duration of each held pose as the model’s prediction for that trial (8 trials per pose in each session; see Training data corpora for details).

##### Architecture

The wrist pose network architecture took as input our custom MPF features of the sEMG signal. The network architecture consisted of two convolutional blocks, with each block consisting of a convolutional layer (dropout=0.1), followed by layer normalization, and a fully-connected layer (ReLU activations). Each layer in the network stack consisted of 128 hidden units and the kernel size of the convolutional layer varied between 1 and 3.

##### Optimization

We trained each network with the AdamW optimizer for a maximum of 400 epochs, with a learning rate of 5e-5 and weight decay of 1e-3. We used a cosine annealing learning rate scheduler, with a minimum learning rate of 1e-8 at 400 epochs. Using the validation loss (cross-entropy loss of the validation data), we performed early stopping when validation performance did not improve for 50 epochs. We evaluated the test performance of the training checkpoint with the lowest validation loss.

#### Discrete Gestures

To train discrete gesture models, we segmented training data from participants into groups of 40, 80, 160, 320, 640, 1280, 2800 and 4800 participants. For each group, we tested the generalization performance of models on offline data from the same set of 100 held-out participants. For validation, another set of held-out users was used; we employed a random set of 16 users for the training groups of size 40 and 80. Subsequently, for larger groups, 10% of the training users were utilized for validation. Each dataset used in training, validation, and testing contained recordings from only a single session per participant. For the largest model, dented with a separate marker on the graph in Fig. 2, we used 4800 training participants and we included multiple sessions of data when available (i.e. many participants collected multiple repeats of the open-loop training protocol). This last point was not included in the fitting procedure for the scaling law, but this model was used in the closed-loop evaluations.

Architecture: The discrete gestures network architecture took as input high-pass filtered sEMG signal. The high-pass filtering was applied with a cutoff of 40Hz with the choice of cutoff determined empirically and primarily motivated by the need to remove motion artifacts (De Luca et al. 2010). The network architecture consisted of a convolutional layer with a stride of 10, reducing the input frequency from 2kHz to 200Hz. This was followed by a layer norm layer, three LSTM layers, another layer norm layer and a linear readout layer corresponding with nine classes, corresponding to the number of gestures. For the smaller model, the dimensions of the convolutional layer and the number of hidden units in the recurrent blocks were set to 128. For the larger model, they were set to 512.

Optimization: Networks were trained using the AdamW optimizer. To mitigate divergence during training, gradient clipping was applied throughout. We further utilized a learning rate scheduler that decayed the learning rate by a factor of 0.5 after a fixed number of epochs (i.e., every 25 epochs). The learning rate for the smaller model was initially set to 1e-3, whereas for the larger model, it was set to 5e-4. Training was done using a binary cross-entropy loss. Each model was trained for a consistent wall clock duration. Final checkpoints were selected based on the model that yielded the highest validation score, defined as classification accuracy.

#### Handwriting

To train handwriting models, we employed the CTC loss as described by (Graves et al. 2006). Notably, we used characters instead of phonemes for this purpose. The characters predicted included all lower-case letters [`a-z`], numbers [`0-9`], [`space`], and punctuation marks [`,.?’!`]. Additionally, we integrated a greedy implementation of the FastEmit regularization technique (Yu et al. 2020). This regularization approach effectively reduced the streaming latency of our models by penalizing sequences of ‘blank’ outputs.

Nine training corpora were generated, each containing a varying number of participants ranging from 25 to 6527 in a geometric sequence (excluding the last point). Each corpus was a superset of the previous corpus’s participants, ensuring that participants in the 25-participant corpus are also present in the 50-participant and 100-participant corpora, and so on. Participants were uniformly sampled without replacement from the entire corpus, preserving the distribution of data quantity per participant found in the full corpus. We used 100 held-out participants to create our evaluation corpora, which remained constant throughout our investigation. The validation corpus comprised data from 50 participants and was utilized for early stopping during model training. The test corpus contained data from 50 participants and served for the final evaluation of each handwriting model’s generalization performance. We also used a subset of these 50 test participants for our personalization corpus (see Personalization experiments).

Two primary data augmentation strategies were employed. The first involved SpecAugment (Park et al. 2019), which applies time- and frequency-aligned masks to spectral features during training. The second strategy involved rotational augmentation, randomly rotating all channels by either −1, 0, or +1 position uniformly. This meant that channel signals were shifted one channel to the left, remained unshifted, or were all shifted to the right, respectively.

For evaluating the model’s offline performance for each user, we used the character error rate aggregated over all prompts collected for that user:

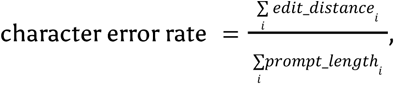

where *edit_distance_i_*is the Levenshtein distance between the prompt and the model output for prompt *i* and *prompt_length_i_* is the length of the prompt.

##### Architecture

The handwriting network architecture took our custom MPF features of the sEMG signal as input. Initially, these features underwent a rotational-invariance module, which comprised a fully-connected layer with LeakyReLU activation. This module was applied to sEMG channels that were discretely rotated by +1, 0, and −1 channels. The resulting outputs were then averaged over the rotation process. It is important to note that this rotation was performed on top of the channel rotation augmentation. The signal was then passed through a conformer (Gulati et al. 2020) architecture consisting of 15 layers. Each layer encompassed 4 attention heads and employed a time-convolutional kernel with a size of 8. Throughout the conformer layer convolutional blocks, a stride of 1 was employed, except for layers 5 and 10 where the stride was set to 2. To ensure the model functioned in a streaming manner, a modified conformer architecture was used. This adaptation is similar to the approach outlined in (Beltagy, Peters, and Cohan 2020), but with adjustments to ensure causality. Specifically, self-attention is solely applied to a fixed local window situated directly before the current time step. In our networks, the size of this attention window was 16 for the initial 10 conformer layers and then decreased to 8 for the subsequent 5 layers. Finally, the outputs from the conformer blocks were subjected to average pooling across channels. They were then passed through a linear layer, which projected the output to match the size of the character dictionary. A softmax function was applied thereafter. During decoding, the model’s best estimate at each output time step was greedily followed, and repeating characters in the prediction were removed to reduce the output.

In our investigation, we explored various trainable model parameter counts. We manipulated the parameter count of our models by adjusting the feed-forward dimension and input dimension within our conformer architecture. Importantly, we upheld a consistent 1:2 ratio between the input dimension and the feed-forward dimension in the conformer blocks.

##### Optimization

The training of our conformer architecture was executed using AdamW as the optimization algorithm. This process spanned a maximum of 200 epochs and involved a learning rate set at 6e-4 for the 1M parameter model and 3e-4 for the 60M parameter model, both with a weight decay of 5e-2. A cosine annealing learning rate schedule was implemented, featuring a warm-up period lasting 1500 steps and a minimum learning rate of 0. Our chosen batch size was 512, wherein each sample within the batch represented a prompt that was zero-padded to match the length of the longest prompt within that batch. To prevent gradient explosion, we applied gradient clipping with a norm threshold of 0.1 throughout the training process. It is worth noting that these parameters were determined based on insights gained from prior experiments and hyperparameter explorations. Lastly, we assessed the test performance of the training checkpoint corresponding to the lowest validation character error rate.

### Online evaluation

#### Study participants

For the wrist pose control and handwriting tasks, we used, as closed-loop study participants, *N=20* Meta employees with less than 1 hour of self-reported previous experience. 18/20 participants reported no previous experience whatsoever with either of these interactions. For the discrete gestures task, *N*=24 naive participants were recruited with no prior knowledge or experience with sEMG wristbands. Demographic information about these participants is shown in Extended Data Fig. 7d-g.

#### Task structure

All closed loop experiments were structured into three blocks: practice block, evaluation block #1, and evaluation block #2. During the practice block, participants were explicitly instructed to explore performing the required gestures/movements in different ways in order to understand how to best perform the task. Explicit coaching was provided during this block to guide participants towards behaviors that were known to be typically well suited for the given sEMG decoder (e.g. “do not press too hard when doing an index/middle press”). During the evaluation blocks, participants were instructed to be as performant as possible, and coaching was only provided when necessary if the participant was stuck on a given trial.

#### Wrist pose control

To evaluate continuous closed-loop control with the wrist pose decoder, we converted the decoder outputs into smoothed velocities to provide velocity-based control similar to a joystick. Specifically, the decoder output probabilities, *p_t_^extension^*, *p_t_^flexion^*, *p_t_^neutral^*, control horizontal cursor positions, *x_t_*, as follows:

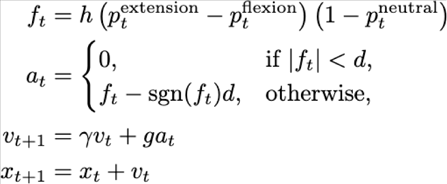

with gain *g=0.2*, dead zone size *d=0.07*, damping factor γ*=0.7*, and *h = 1* if the sEMG wristband is on the right hand (so that wrist flexion/extension maps to left/right, respectively) and *-1* if it is on the left hand (so that wrist flexion/extension maps to right/left).

We evaluated cursor-control performance using a 1D cursor control task, in which participants were prompted to move the cursor to one of 10 equally-sized rectangular targets presented on a horizontal grid (see Fig. 3a, Extended Data Fig. 6a, Supplementary Video 1). At each trial, one of the 10 targets was highlighted, and the participant was instructed to move the cursor towards that target. The target was acquired when the cursor remained within the target for 500ms. Once a target was acquired, the rectangular target disappeared, and one of the remaining 10 targets was prompted, initiating the next trial. Once all 10 targets were acquired, a new set of 10 targets was presented, and each one was prompted in a random sequence. This was repeated 5 times in each block, for a total of 50 trials per block. The cursor position was continually decoded from sEMG throughout the session and never reset between trials or blocks. The cursor workspace was bounded at the outer edges of the leftmost and rightmost targets.

All trials with an initial cursor distance less than 10% of the target width were discarded (56/3000 trials). These occurred whenever, by chance, the next prompted target happened to be immediately next to the current cursor position. Such serendipitous trials had abnormally low acquisition times so were discarded as outliers. Fig. 3d shows the mean acquisition time over all non-discarded trails in each block, for each participant. The 500ms hold time is not included in the acquisition time. Note that this average is over trials with varying starting distances from the target. In Extended Data Fig. 7, we further examine performance in this task using Fitts’s law throughput (Fitts 1954), which accounts for trial-to-trial differences in reach distances and has been previously used in HCI (MacKenzie 1992) and BCI settings (Gilja et al. 2012).

An additional measure we used to quantify performance was dial-in time (Fig. 3e), which is a measure of precise control around the target, adapted from the BCI literature (Fan et al. 2014). Dial-in time was measured as the time from the first target entry to the last target entry, not including the 500ms target hold time. Fig. 3e shows the mean dial-in time over all non-discarded trials in which the cursor prematurely exited the target before completing the 500ms hold time (i.e. trials in which the dial-in time was greater than 0).

#### Discrete gestures

To evaluate the discrete gesture decoder in closed loop, we used a discrete grid navigation task in which each of the thumb swipes (left/right/up/down) was used to move a yellow circular character, named Chomper, along a discrete grid (see Fig. 3b, Extended Data Fig. 5b, Supplementary Video 2). Movements were prompted with a series of targets indicating the direction in which Chomper should move, and every few steps the participant was prompted to perform one of the three “activation” gestures: thumb tap, index hold or middle hold. Index/middle holds were defined as a press followed by a release at least 500ms later. If the release followed the press less than 500ms later, this was classified as an “early release” error. If the model detected a thumb swipe in the wrong direction, Chomper would move in the detected direction and the participant would thus be prompted to swipe in the opposite direction to move Chomper back to its previous position. The total number of prompted gestures in each trial could thus vary depending on how many times the wrong thumb swipe direction was detected. Incorrect activation detections would be indicated to the participant, but would not alter Chomper’s position. Each trial consisted of a randomly sampled sequence of 8 thumb swipes and 5 activations, with 10 trials in each block. Participants were explicitly instructed to favor accuracy over speed when performing the task.

Completion rate (Fig. 3g) was defined as the time required to complete a trial divided by the minimum number of discrete gestures required to complete a trial (8 thumb swipes + 5 activations = 13 gestures). Mistakenly making additional gestures that were counterproductive to completing the trial added to the time required, but did not increase this number of required gestures. To calculate the confusion matrix for each participant, we counted the number of times each gesture was detected when a given gesture was expected. To get a proportion, we then divided this by the total number of gestures executed when that given gesture was expected. Fig. 3h shows the average confusion matrix across all participants, using the trials in the two evaluation blocks only. The first hit probability (Fig. 3f) was calculated by taking the proportion of prompted gestures in which the first executed gesture was the expected one. For both the first hit probability and the confusion matrix metrics, we included the 13 prompted gestures in each trial as well as any additional prompted thumb swipes resulting from swiping in the wrong direction.

Note that in order to complete the discrete gestures task, the participant was required to perform all gestures correctly. So before this task began, all participants were screened to confirm that each gesture worked for them; however, no participants had prohibitive issues with any gesture.

#### Handwriting

To evaluate the handwritten character decoder in a closed loop, we used a handwriting task in which participants were prompted to handwrite 5-word phrases sampled from the Mackenzie corpus (Mackenzie and Soukoreff 2001). Characters ([`a-z`], [`0-9`], [`space`], [`,.?’!`]) were decoded online with the decoder and displayed to the participant in real time (see Fig. 3c, Extended Data Fig. 5c, Supplementary Video 3), and participants were instructed to ensure that the decoded phrase was understandable before submitting it and moving on to the next trial. If the participant produced any incorrect characters, they could use the backspace key on the keyboard to erase errors and then re-write them. Trials were completed when participants made their best attempt to write the prompted phrase and then submitted the written text by pressing a key on the computer keyboard using their non-dominant hand. Each block consisted of 10 such trials.

In our analysis, we report the aggregate character error rate (CER) and adjusted words per minute (aWPM) over all prompted characters in each block, calculated as in (Palin et al. 2019). For each block, the character error rate was aggregated over all 10 prompts as follows:

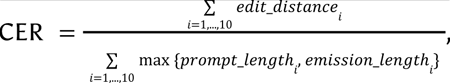

where *edit_distance_i_* is the Levenshtein distance between the prompt and the model output submitted by the user in trial *i*, *prompt_length_i_* is the length of the prompt, and *output_length_i_* is the length of the model output. The maximum between these two is used in the denominator in order to ensure the aggregate CER is between 0 and 1 for the adjusted WPM metric to be meaningful.

### Discrete gesture detection model investigation

#### Network convolutional filter analysis

To examine the initial Conv1d layer of the trained discrete gesture decoder, we first measured various spatiotemporal properties of each of the Conv1d filter weights. We measured the root mean square power in each channel and identified the channel with maximum power. We then measured the frequency response of this max channel using a discrete Fourier transform and identified the peak frequency with strongest magnitude response. We measured the bandwidth of the frequency response as the range of contiguous frequencies around this peak that had a magnitude response within 50% of the peak. We additionally counted how many channels had root mean square power within 50% of the max channel. The distributions of these metrics across all Conv1d filters are shown in Extended Data Fig. 8. We next identified the set of Conv1d filters that fell within the interquartile range of these three metrics (peak frequency, bandwidth, number of active channels), and randomly selected six filters with different peak channels. These are the representative examples shown in Fig. 4b, 4d, 4e. The six putative MUAPs shown in Fig. 4c were extracted using the procedure described in Online Methods, and then the raw EMG signal in the central 10ms of each snippet was high-pass filtered with the same preprocessing procedure applied to the discrete gestures model training data (Online Methods). This allowed a direct comparison with the 10ms convolutional filters trained on data preprocessed in this way. The same procedure for measuring RMS power and frequency response was applied to the six putative MUAPs after this preprocessing to obtain the curves shown in Fig. 4d, 4e.

#### Discrete gesture detection network LSTM representation analysis

To examine the LSTM representations of the trained discrete gesture decoder, we used recordings from 3 different sessions from each of 50 randomly selected users from the test set. From each of these recording sessions, we randomly sampled forty 500ms sEMG snippets ending at labels for each gesture class (after label timing alignment, see Online Methods), for a total of 40 x 9 = 360 sEMG snippets per session. We then passed each of these snippets through the trained discrete gesture decoder, with the LSTM state initialized to zeros, to obtain vector representations, *X* ∈ *R*^512^, of each snippet. These are the vectors whose PC projections are plotted in Fig. 4f-h, in each case colored by a different property. Gesture-evoked sEMG power was measured as the root mean square of the last 100ms of each sEMG snippet. For each participant and gesture, this was then binned into 20 bins with a matched number of snippets, dividing the EMG power into the categories plotted in Fig. 4h.

To quantify the structure in these representations, we used the proportion of variance in LSTM representations accounted by a given variable, ξ:

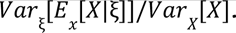

The numerator is the variance in the mean representation for each category of ξ, and the denominator is the total variance of the representations. In each case, variance is calculated as the trace of the covariance of the representations. For the discrete gesture identity and participant identity analysis, we divide the 50 participants into 10 non-overlapping sets of 5 participants and calculate the proportion of variance separately for each set. The curves in Fig. 4i show the mean and 95% confidence interval over these 10 sets. For the band placement and gesture-evoked EMG power curves, the proportion is calculated separately for each of the 50 participants, and the mean and 95% confidence interval over participants is shown. For this analysis, the EMG power was binned as indicated above but into only 3 bins (low/medium/high) rather than 20.

### Personalization experiments

We studied the personalization of handwriting models with 40 participants from the test corpus that were held out from the 6527 participants in the pre-training corpus. For each participant, we further trained, i.e., fine-tuned, a chosen generic handwriting model on a fixed budget of data solely taken from that participant’s sessions. The resulting personalized model was then evaluated on held-out sessions of data from the same participant it was personalized on. We considered personalization data budgets of 1, 2, 5, 10 and 20 minutes. We repeated this process for each of our 40 participants and reported the population average of the personalized model performance.

#### Data

We created a training and testing set for each of our 40 personalization participants by holding out three sessions for the test set and keeping the rest of the sessions in the training set. The three held out testing sessions were chosen such that there was one session recorded in each of the three postures (seated writing on a surface, seated writing on their leg, and standing writing on their leg). The training set, obtained by gathering the rest of the participant’s sessions, was subsetted in order to obtain the desired number of minutes of labeled recording. The subsetting was done through random uniform sampling of the prompts from all of the sessions in the training set. Each subset of the full training set was a superset of the precedent data budget size, ensuring that the prompts in the 1-minute budget were also present in the 2-minute and 5-minute budget, and so on.

#### Optimization

The optimization details closely resemble the procedure followed for generic training (See Generic Model training) with a few differences, which were that we used a cosine annealing learning rate schedule without warmup. Also, we varied the fine-tuning learning rate as a function of the number of pre-training participants used to pre-train the upstream generic model, such that: *LR*(*N*) = 1. 24 * 10^−5^ * *N*^−0.42^, with N being the number of pre-training participants. The learning rate relationship with generic pre-training participants was found through a grid learning rate sweeps for the models pre-trained on 25, 400, and 6527 participants, then fitting a power law to the population average performance minima found. We did not use weight decay during fine-tuning. We fine-tuned the model for 300 epochs, at a batch size of 256, with no early stopping such that the training is always 300 epochs.

#### Statistics

In Fig. 5e, we find negative transfer of personalized models across participants. In order to characterize each participant’s performance on other fine-tuned models, we first compute the mean of each row without the diagonal. We then compute the median of the means along with the standard error of the median (s.e.m.). This is compared with the median of the diagonal values.

In Extended Data Fig. 9, we added early stopping to the personalization procedure to disambiguate the contribution of increased personalized data budget per user from an increase in the number of fine-tuning iterations. We find very similar results with (Extended Data Fig. 9) and without (Fig. 5 of the main text) early stopping, except a few of the best performing users exhibit regressions from personalization without early stopping. This verifies that the benefits from including more personalization data are not due to an increase in training iterations. Note that in practice early stopping would require additional data from the participant to use for validation. Here we used the test set for early stopping, so the results in Extended Data Fig. 9 should be considered validation numbers.

### Fitting procedures

#### Generic performance fitting

##### Fitted function

In Fig. 2d-f, we show the fits of the generic error scaling with the number of training participants. The fits follow a functional form taken from the large language model literature (Hoffmann et al. 2022), where the error is a function of both model size (*D*, in number of parameters) and data quantity (*N*, in number of participants):

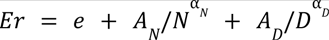

where all fitted parameters are positively bounded. It is generally understood that the *e* term in this equation is the irreducible error of the task and the the second and third terms both contribute to the error reduction as *N* and *D* are increased, respectively. Note that there exists diminishing return regimes if either *N* or *D* are increased individually, as the other term fixes the asymptotic error floor.

##### Fitting procedure

A single set of parameters fit all of the observed points in each graph, with the exception of the heterogeneous data point in the discrete gestures experiments which we keep held out because of its training corpus distinction with the rest of the points. The fitted parameters were obtained by minimizing the Mean Squared Logarithmic Error (MSLE) using the L-BFGS-B optimization algorithm (Byrd et al. 1995) along with 200 iterations of the basin hopping strategy (Olson et al. 2012). The initial guess and the bounds for the fitted parameters are shown in Table 1.

**Table 1.** Parameters for power law fits in Fig. 3.

| Parameter | $e$ | $A_N$ | $\alpha_N$ | $A_D$ | $\alpha_D$ |
| --- | --- | --- | --- | --- | --- |
| Initial guess | 0 | 1 | 1 | 1 | 1 |
| Bounds | $[0, \infty)$ | $[0, \infty)$ | $[0, \infty)$ | $[0, \infty)$ | $[0, \infty)$ |
| Wrist pose | 2.6 | 7.7 | 0.33 | 13.5 | 0.12 |
| Discrete gestures | 4.2 | 12.3 | 0.33 | 0.1 | 2.68 |
| Handwriting | 2.1 | 26.1 | 0.36 | 3.6 | 2.65 |

### Personalization experiment fitting

#### Fitted function

In Fig. 5b, we show the fits of the 60M parameter model error rate as a function of the number of pre-training participants for the generic model and each personalization data budget. The fits follow a functional follow a simple power law fit with the pre-training data quantity (*N*, in number of participants), such that:

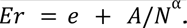

We did not take into consideration the contribution from model size, since we are only fitting observations from a single model size (hence the error from finite model size is absorbed into *e*).

#### Fitting procedure

The fitted parameters for each personalization data budget were obtained by minimizing the Mean Squared Logarithmic Error (MSLE) using the L-BFGS-B optimization algorithm (Byrd et al. 1995) along with 200 iterations of the basin hopping strategy (Olson et al. 2012). The initial guess and the bounds for the fitted parameters are shown in Table 2.

**Table 2.**
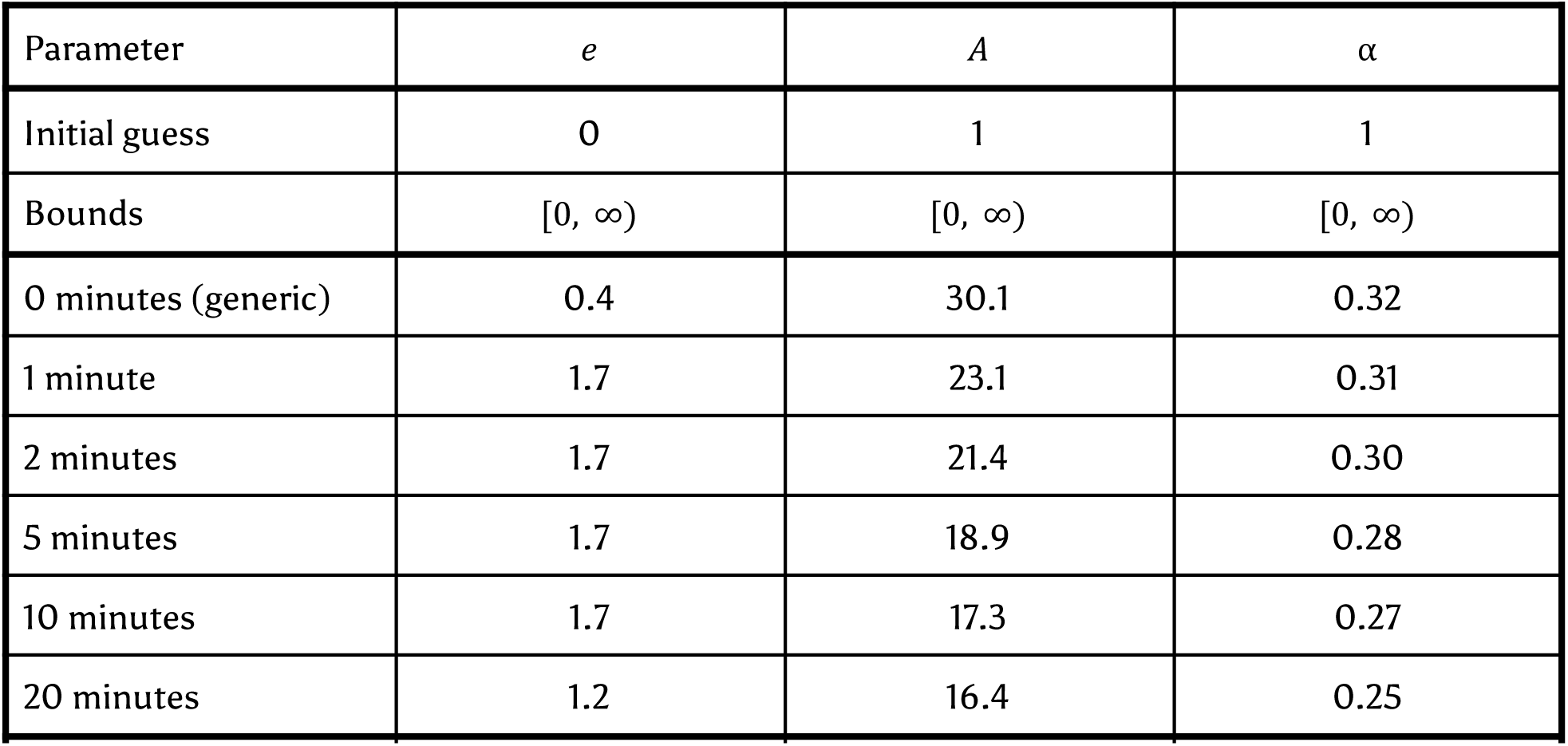
Parameters for power law fits in Fig. 5.

| Parameter | $e$ | $A$ | $\alpha$ |
| --- | --- | --- | --- |
| Initial guess | 0 | 1 | 1 |
| Bounds | $[0, \infty)$ | $[0, \infty)$ | $[0, \infty)$ |
| 0 minutes (generic) | 0.4 | 30.1 | 0.32 |
| 1 minute | 1.7 | 23.1 | 0.31 |
| 2 minutes | 1.7 | 21.4 | 0.30 |
| 5 minutes | 1.7 | 18.9 | 0.28 |
| 10 minutes | 1.7 | 17.3 | 0.27 |
| 20 minutes | 1.2 | 16.4 | 0.25 |

### Personalization equivalence calculations

#### Relative increase calculation

To determine the equivalent pre-training participant budget needed to match a given personalization performance, we need a continuous estimate of generic model performance as a function of the number of pre-training participants. For this, we used logspace piecewise linear interpolation of the generic performance values, which we denote by *f_generic_*(*N*). Given the number of pre-training participants, N, and personalization minutes, m, personalized models have an observed character error rate given by CER(N, m). To find the equivalent additional pre-training participants Δ*N* needed to match performance between generic and personalized models we set *f_generic_*(*N* + Δ*N*) = *CER*(*N*, *m*) and solve for Δ*N* using the Newton Conjugate-Gradient method. This gives the points in Fig. 5(d). Overlaid on the plot as dotted lines, we used the power law fit of the points corresponding to each number of personalization minutes in Fig. 5(b) to infer continuous curves of equivalent fold-increase in pre-training data required using the approach described above.

## Supplemental figures

**Extended Data Fig. 1.**
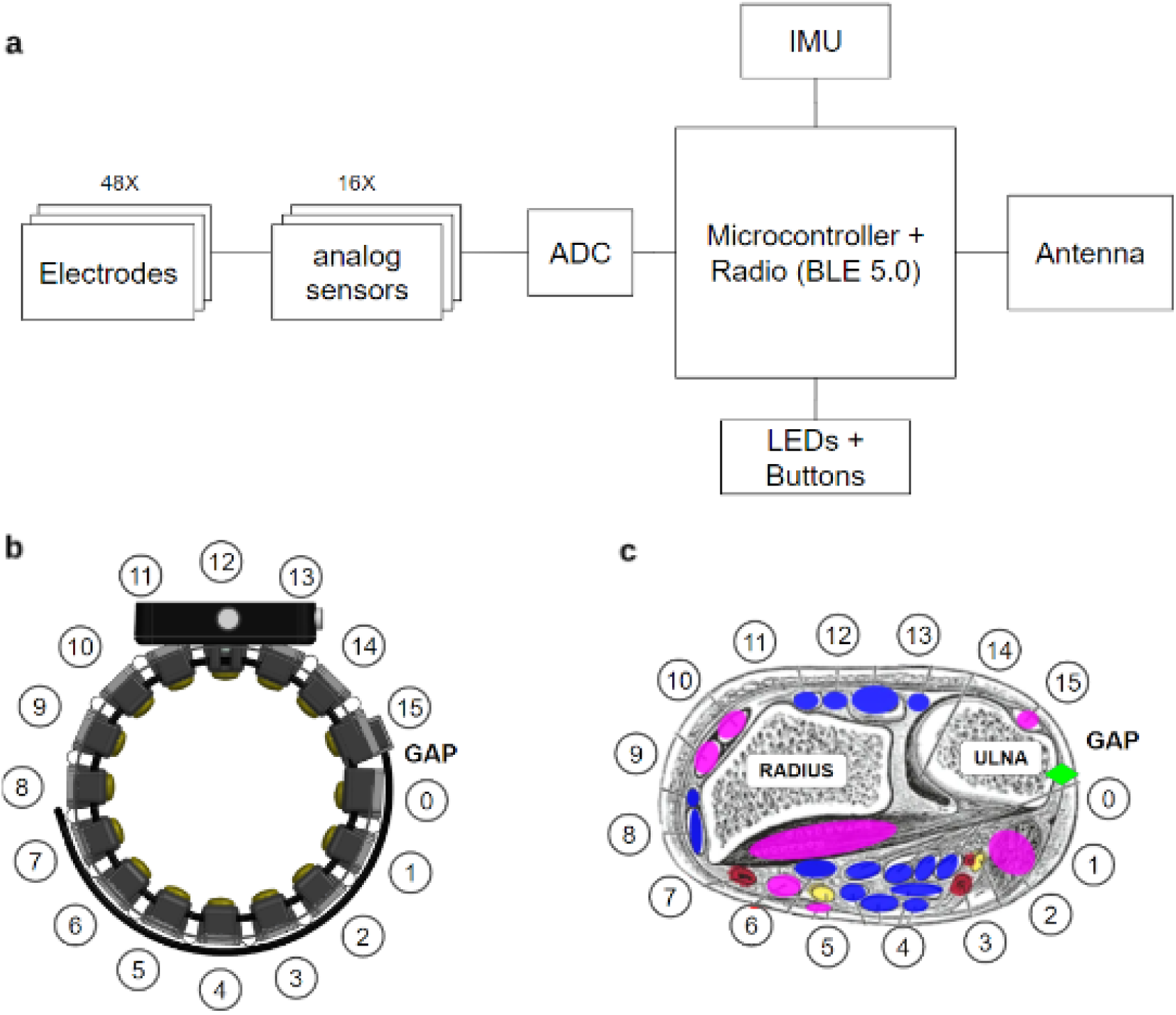
Schematic and anatomical interfacing of sEMG Research Device. **a**, The sEMG Research Device electrical system architecture. The sEMG-RD uses 48 pogo-pin style round electrodes in order to provide good comfort and contact quality. The 48 channels are configured into 16 bipolar channels arranged proximo-distally, with the remainder electrodes serving as either shield or ground. Each electrode is 6.5 mm in diameter (gold plated brass). For each differential sensing channel (16 in total), center-to-center spacing between paired sensing electrodes is 20 mm. The sEMG-RD has low noise analog sensors with input-referred RMS noise of 2.46 µVrms, measured during benchtop testing with differential inputs shorted to their mid-point voltage. With analog sensors’ nominal gain value of 190 and Analog to Digital Converter’s (ADC) full-scale range 2.5 V, the sEMG-RD offers a dynamic range of approximately 65.5 dB. Each channel is sampled at 2000 Hz. The Inertial Measurement Unit (IMU) functional block includes sensors of 3-axis accelerometer, 3-axis gyroscope, and 3-axis magnetometer sampled at 100 Hz. We note that the IMU was not utilized for any online or offline experiments described in this manuscript. The microcontroller facilitates the transfer of unprocessed data from all ADCs and IMU directly to the bluetooth radio. No skin preparation or gels are needed for using the sEMG-RD, because its analog sensors have very high input-impedance – approximately 10 pF capacitance in parallel with 10 TOhm resistance—providing excellent signal robustness against large variations of electrode-skin impedance among the population. **b**, The mechanical architecture consists of a kinematic chain with flexible joints connecting 16 pods that house the sprung electrodes that comprise the sEMG channels. This enables broad population coverage in maintaining consistent quality contact between the dry electrode and skin. Since each differential sensing channel is placed along the proximal-distal direction, the device is able to maintain symmetry with respect to wrist anatomy and provide generalizability across right and left hands, as long as the wearer keeps the gap location on the ulna side. **c**, Anatomical depiction of electrode locations relative to relevant muscle and skeletal landmarks, adapted from a public domain image (Gray 1918). Pink overlays cover muscles that predominantly control the wrist, blue overlays cover muscles less involved in wrist control, red overlays cover blood vessels and yellow overlays cover nerves. The green diamond indicates the position of the electrode gap. Note the gap that arises between channels 0 and 15, due to variation in wrist circumference and elasticity between compartments, is aligned with the region of the wrist where the ulna is located.

**Extended Data Fig. 2.**
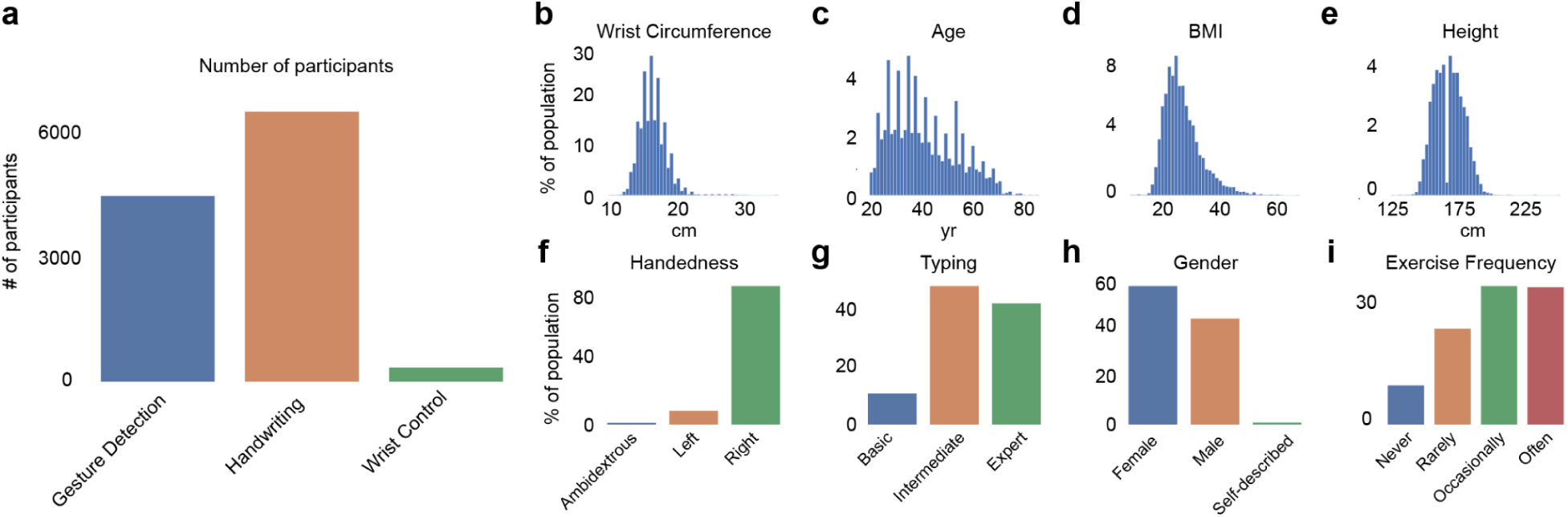
Anthropometric and demographic features of sEMG datasets. **a**, The number of participants in each corpus. **b-e**, Histograms of anthropometric characteristics of participants: (**b**) wrist circumference, (**c**) self-reported age, (**d**) self-reported BMI, and (**e**) self-reported height. The irregularity in the histogram of self-reported age is likely due to participants rounding their age to nearby values. We measured wrist circumferences with a standard measuring tape at the wrist just below the ulna bone where the participants are expected to don the band. Values outside of the range of 10-30 cm were truncated. We calculated BMI as the weight (in kilograms) divided by height (in meters) squared. **f-i**, Distributions of the demographic characteristics across all participants: (**f**) dominant handedness, (**g**) self-reported proficiency at typing on a computer keyboard, (**h**) self-reported gender, and (**i**) arm exercise frequency, chosen from one of the following options: Never (never), Less than once per week (rarely), 1-2 times per week (occasionally), more than twice per week (often).

**Extended Data Fig. 3.**
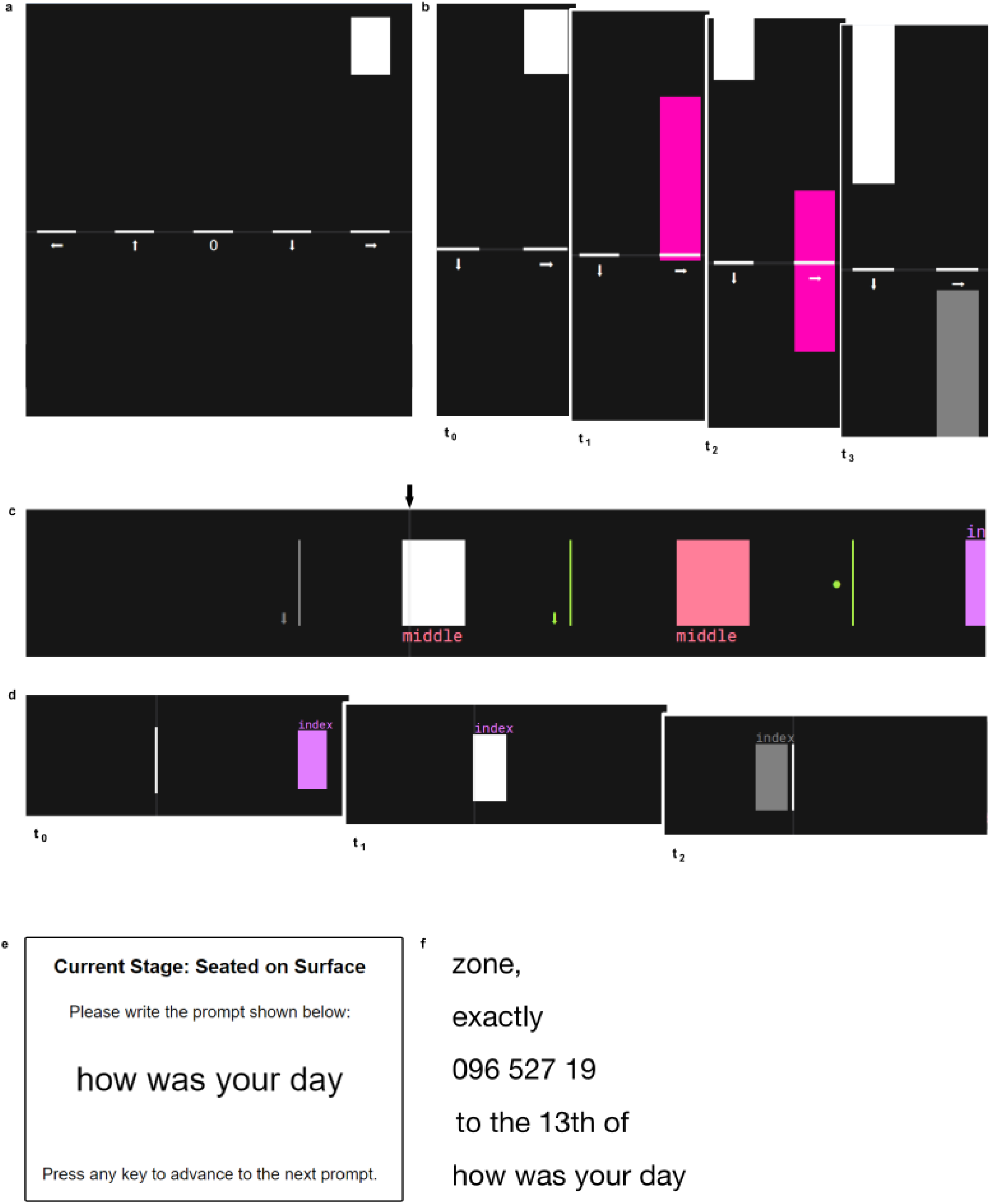
Examples of open-loop prompting for the three tasks. **a**, Example prompter frame used to train the model for the wrist prediction task. Vertical bars move from the top of the screen to the bottom. A series of indicator lines labeled with different directions were shown at the bottom-middle of the screen. The participant is instructed to perform the appropriate pose when a descending bar reaches the indicator line, e.g. extend/flex their wrist horizontally or deviate their wrist vertically, following the direction marked on the indicator line overlapping with the bar. ‘0’ corresponds to a neutral wrist pose. The pose is held until the entire bar has passed the indicator line. **b**, Time series of example prompter frames for the wrist pose task. At t_0_ a bar appeared above the indicator line for ‘wrist right’. At t_1_ the bar reached the indicator and the bar changed color to pink, indicating that the participant should move their wrist to the right. At t_2_ the color was maintained, indicating that the participant should continue to hold the wrist posture, and a second descending bar appeared above the ‘wrist down’ indicator line. At t_3_ the completed ‘wrist right’ bar has changed color to gray, indicating that the participant should return their wrist to a neutral posture, and the ‘wrist down’ bar continues to descend towards the indicator line. c, Example of prompting for the discrete gesture recognition task. A series of gestures to be performed are depicted, with colors and labels corresponding to the gesture type. Gestures are separated by blank space in which no gesture was to be performed. These prompts scroll similarly to the wrist pose prompter, though leftwards instead of downwards. When the prompt reaches the indicator line, gestures should be initiated – either instantaneous gestures such as finger pinches or thumb swipes that are depicted as single lines or held gestures such as index and middle holds that are depicted as solid bars. Held gestures should be released when the indicator line reaches the end of the rectangle. Gestures that have already been prompted are shown in gray. **d**, Detailed example of prompting during holds. At t_0_ an index hold gesture prompt appeared on the right side of the screen, with the time indicator line in white. At t_1_ the gesture prompt reached the time indicator, and the hold prompt changed color to indicate the hold should be performed by the participant. At t_2_ the hold was no longer selected by the indicator bar and turned gray, indicating that the participant should release the hold. **e**, Example prompter shown during the handwriting task. The screen instructed the participant to write “how was your day” with their hand on the surface of the table, while seated. **f**, During the experimental session, different prompts, including numbers and punctuation, were shown, ranging from single characters to full sentences. Besides writing on a desk surface, the participant was also asked to perform handwriting on their leg while standing and on their leg while seated.

**Extended Data Fig. 4.**
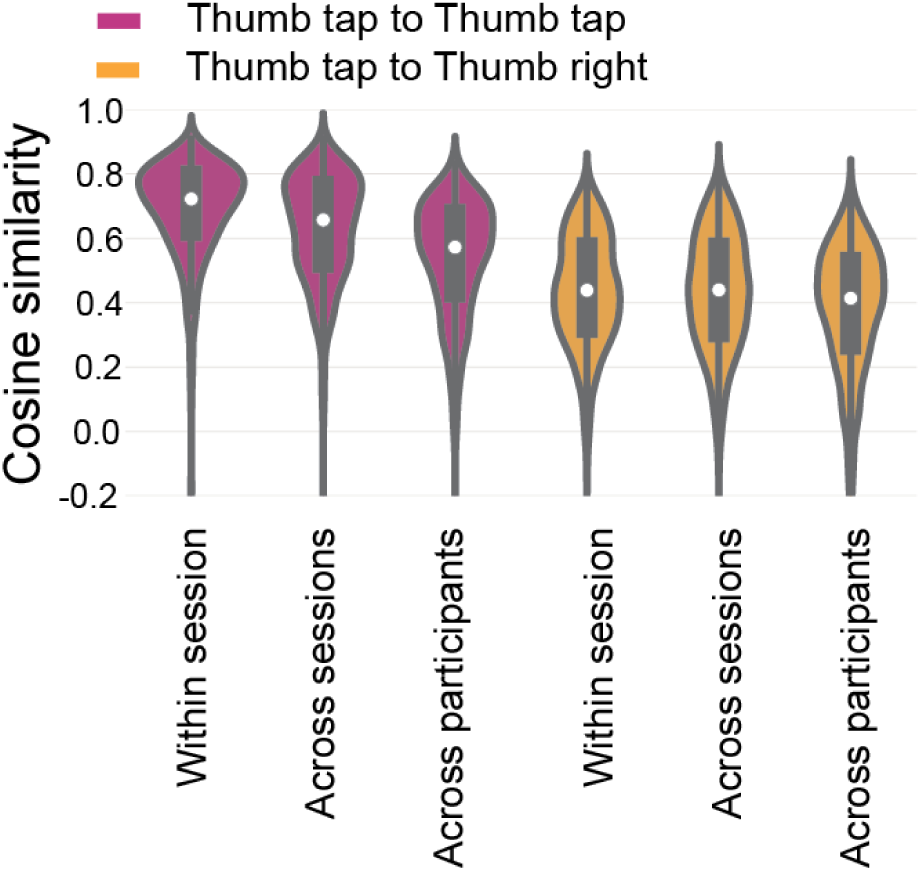
Distribution of sEMG event similarities across participants and gestures. *Purple*: cosine similarity between individual sEMG activations of a given gesture and the sEMG template (event-triggered average) for that gesture. From left to right: cosine similarities are plotted for all events within a single session (single band placement), across sessions of a single participant, or across all participants from Fig 2a. While similarity was relatively high within a single band placement, sEMG activations became progressively more distinct across different band placements and individuals. *Orange*: same, except for the cosine similarity of one gesture compared to the template for a distinct gesture. These were lower than similarity within the same gesture, irrespective of whether the grouping was done over a single band placement or across the population. Differences shown across sessions, participants and gestures are representative for all gestures and pairs of gestures.

**Extended Data Fig. 5.**
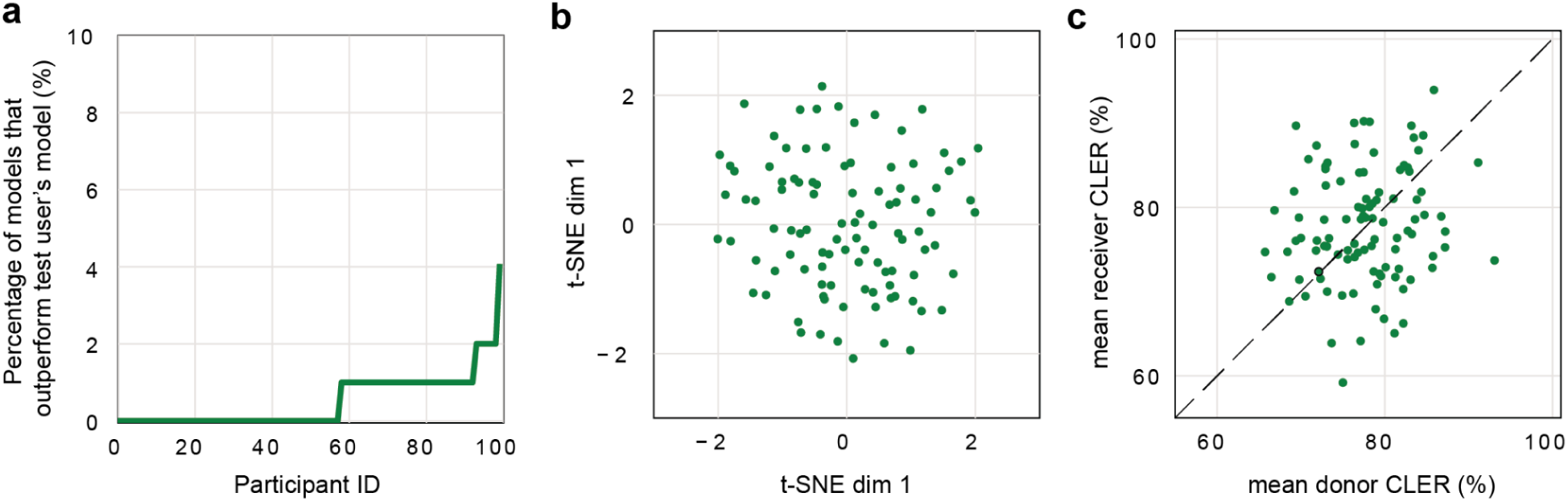
Lack of overt structure in single-participant sEMG decoder performance. **a**, For each held-out individual, only a small fraction of other single-participant models in the discrete gesture detection task (Fig. 2c,d) outperformed the person’s own model (green line, <5% for any given person, people sorted by fraction of models that outperformed person’s own model). All the results in this figure are based on models trained on 4 sessions. **b**, Qualitative inspection of t-SNE embeddings reveal no prominent similarity structure when the average model transfer CLER is used as a proxy for distance between two participants. **c**, Scatter plot comparing, for each participant, that participant’s model’s average offline performance on other participants’ held-out sessions (donor score, x-axis) compared to the average performance of other participant’s models on that participant’s held out session (receiver score, y-axis). There is not a significant Pearson correlation between the donor and receiver score (*r*=0.13, *p*=0.19). Individual models that show high CLER fail to generalize across the population and thus show very weak generalization capabilities (green dots show scatter plot between donor and receiver score, and dashed gray line shows *x*=*y* line).

**Extended Data Fig. 6.**
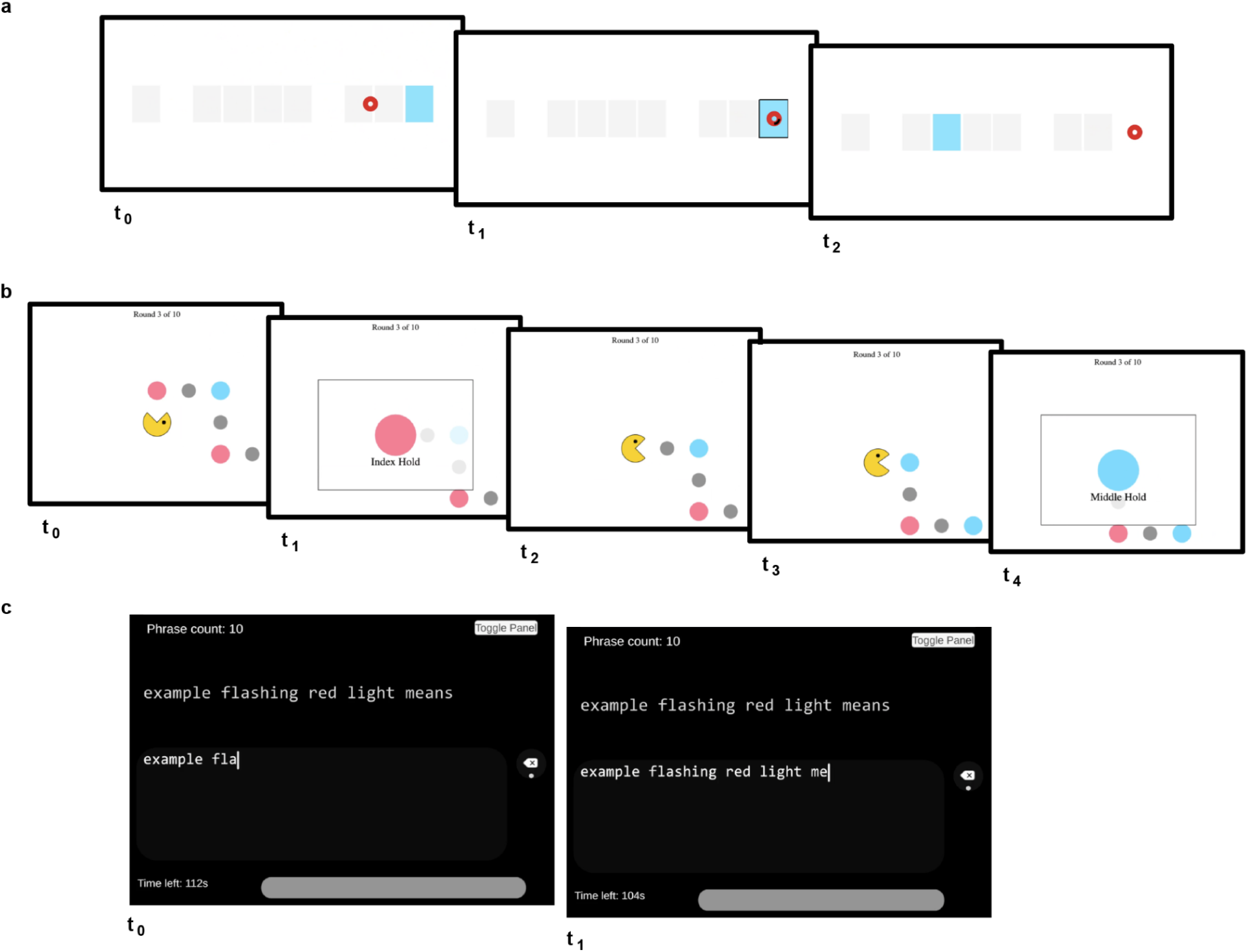
Example screenshots of closed-loop evaluation tasks. **a**, Example trial of 1D horizontal cursor control task, in which the participant was prompted to reach to the rightmost target (in panel labeled t_0_, light blue rectangle). When the cursor (red) landed on the target, a short timer began, marked by the black fill of the cursor (middle panel, t_1_). In this trial, the cursor was held on the target for 500ms to complete the timer, so the target was acquired and therefore disappeared as the next target was prompted (right panel, t_2_). **b**, Example sequence from discrete grid navigation task, in which the participant was prompted to perform (from left to right): thumb swipe up, index hold, thumb swipe right, thumb swipe right, middle hold. **c**, Example handwriting task trial, in which the participant is prompted to write the phrase “example flashing red light means” (top) and the participant’s writing (below).

**Extended Data Fig. 7.**
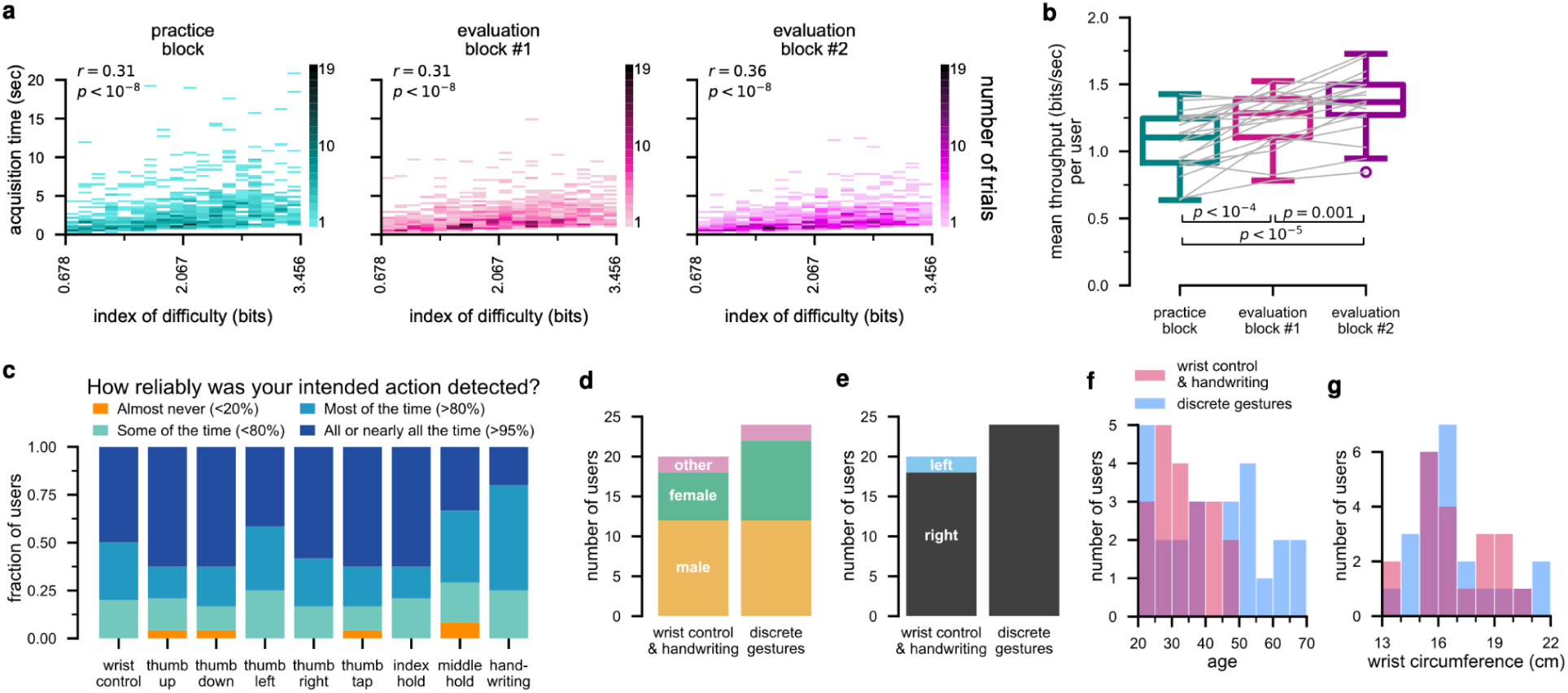
Additional online evaluation metrics. **a**, Distribution of 1D horizontal cursor control task per-trial acquisition times as a function of index of difficulty, log_2_ (1 + *d_t_*/*w*), where *w* is the target width and *d_t_* is the initial target distance in trial *t* (Gilja et al. 2012). The relationship between these two measurements appears to be linear, suggesting behavior in this task obeys Fitts’s law (Fitts 1954). Pearson correlation coefficient and p-value are shown in the upper right hand corner of each panel. **b**, Mean throughput on the 1D horizontal cursor control task. Throughput is defined as the index of difficulty divided by acquisition time. The mean throughput over trials in each block is shown for each participant, following the same conventions as Fig. 3d,e; median in the last block is 1.37 bits/sec. Throughput in each block significantly improved relative to the previous block (*p<.05*, one-tailed Wilcoxon signed-rank test), indicating learning effects consistent with the improvements in acquisition time and dial-in time shown in the main text. **c,** Distribution of subjective impressions about the reliability of each EMG decoding model. At the end of each online evaluation task, participants were asked to respond to a multiple choice question about how reliably their intended action was detected. For the discrete gestures task, they were asked to answer this question separately for each of the thumb swipe directions and “activation” gestures. **d**-**g,** Demographics of *N*=20 participants that performed the wrist pose control and handwriting closed-loop evaluation tasks and the *N*=24 participants that performed the discrete gestures task: (d) self-declared gender, (e) self-declared dominant hand, (f) self-declared age, (g) measured wrist circumference.

**Extended Data Fig. 8.**
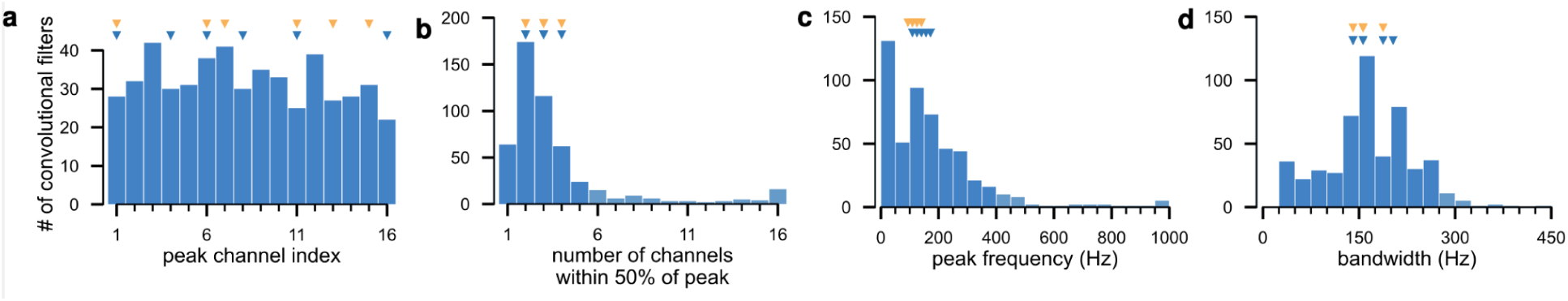
Spatiotemporal properties of all discrete gesture decoder convolutional filters. **a**, Index of channel with max root-mean-square (RMS) power. Here and in all other panels in this figure, the triangles at the top mark the values of the 6 example convolutional filters from Fig. 4b (in blue) and the 6 example putative MUAPs from Fig. 4c (in orange). **b**, Number of channels with RMS power within 50% of the peak channel. **c,** Peak frequency response of the channel with max RMS power. **d,** Bandwidth of the channel with max RMS power (see Online Methods).

**Extended Data Fig. 9.**
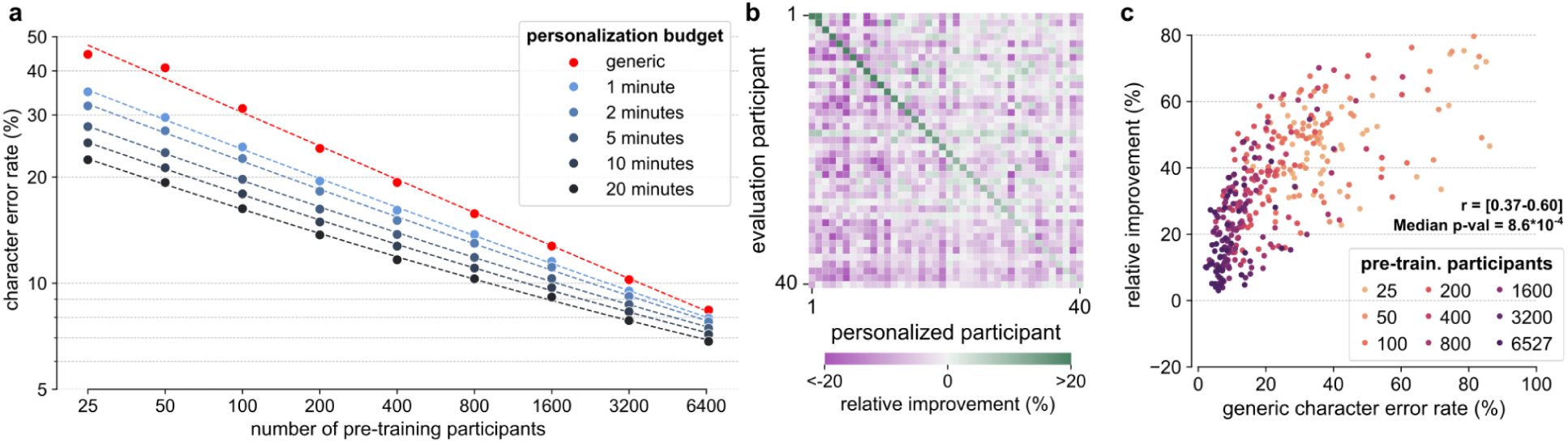
Influence of early stopping during personalization. In this figure, we employ early stopping during personalization to disambiguate the role of more personalization data from increased fine-tuning iterations as well as to mitigate regressions among the best-performing users. Specifically, we used mean CER on held out test data as a selection criteria for epoch-wise early stopping. Aside from early stopping, the setup here is identical to that in Fig. 5 (b, e, f) of the main text. Overall, results are very similar to Fig. 5 of the main text, indicating that the increase in personalization data is the primary driver of improved performance. Regressions among the best-performing users are now absent. Note also that we do not have separate validation and test sets, so these results should be understood as validation performance. **a**, Same as Fig. 5(b) of the main text, except with the inclusion of early stopping during fine-tuning. **b**, Same as Fig. 5(e) of the main text, except with the inclusion of early stopping during fine-tuning. Compared with Fig. 5(e), transfer of personalized models to other participants yields overall smaller regressions likely because early-stopped models remain closer to the pre-trained model. **c**, Same as Fig. 5(f) of the main text, except with the inclusion of early stopping during fine-tuning. Regressions exhibited by a few of the best performing users in Fig. 5(f) are now absent due to early stopping. We show the range of Pearson correlation coefficients for each fit and the median p-value (maximum p=0.020).

## Supplementary Videos

**Supplementary Video 1 | Example session of the 1D horizontal cursor control task with generic wrist pose decoding model**

Example user performing the 1D horizontal cursor control task with the generic wrist pose decoding model evaluated in Fig. 3d-e. In each trial, the participant is prompted to move the cursor to a target and acquire it by holding the cursor within the target region for 500ms. Once that target is acquired, the target disappears and another is prompted. Once all 10 targets are acquired, another 10 identical targets are presented and prompted in random order. Video shows all 50 trials in one task block, where a “trial” is defined as a single target acquisition. Video in the lower right shows the user’s hand during the task.

**Supplementary Video 2 | Example session of the discrete grid navigation task with generic discrete gestures decoding model**

Example user performing the discrete grid navigation task with the generic discrete gesture decoding model evaluated in Fig. 3f-h. In each trial, the participant is instructed to move a character along a sequence of points on a grid by using the four “navigation” gestures (thumb swipe left/right/up/down). Every few steps, a colored point with text prompts the participant to perform an “activation” gesture (thumb tap, index hold, middle hold). Video shows all 10 trials in one task block. Video in the lower right shows the user’s hand during the task.

**Supplementary Video 3 | Example session of the handwriting task with handwriting model**

Example user performing the handwriting task with the generic handwriting decoding model evaluated in Fig. 3i-j. In each trial, the participant is prompted to write a phrase. First part of the video shows all 10 trials in one task block, with the user’s hand on the table as in the experiment reported in the main text. Second part of the video shows the same user writing on their lap, to demonstrate the model’s invariance to hand posture. Video in the lower right shows the user’s hand during the task. While we instructed users to make corrections to decoded text that was not understandable, in this particular session the user did not have to make any such corrections.

**Supplementary Video 4 | High-level explanatory video of technology with examples**

Overview of the technology. A participant can easily don the sEMG-RD, which can detect rich surface electromyogram signals that accompany movement. sEMG decoding models enable multiple human-computer interactions in novel participants, for example 1D continuous control, discrete gesture recognition, and handwriting. Examples of the three tasks in the second half of the video are taken from Supplementary Video 1-3. Example video of handwriting to produce the phrase ‘hello world!’ at the start of the video was made using the generic handwriting decoding model evaluated in Fig. 3i-j.

